# Regulation of the circadian clock in *C. elegans* by clock gene homologs *kin-20* and *lin-42*

**DOI:** 10.1101/2023.04.13.536481

**Authors:** Melisa L. Lamberti, Rebecca K. Spangler, Victoria Cerdeira, Myriam Ares, Lise Rivollet, Guinevere E. Ashley, Andrea Ramos Coronado, Sarvind Tripathi, Ignacio Spiousas, Jordan D. Ward, Carrie L. Partch, Claire Y. Bénard, M. Eugenia Goya, Diego A. Golombek

## Abstract

Circadian rhythms are endogenous oscillations present in nearly all organisms from prokaryotes to humans, allowing them to adapt to cyclical environments close to 24 hours. Circadian rhythms are regulated by a central clock, which is based on a transcription-translation feedback loop. One important protein in the central loop in metazoan clocks is PERIOD, which is regulated in part by Casein kinase 1*ε/δ* (CK1*ε/δ*) phosphorylation. In the nematode *Caenorhabditis elegans*, *period* and *casein kinase 1ε/δ* are conserved as *lin-42* and *kin-20*, respectively. Here we studied the involvement of *lin-42* and *kin-20* in circadian rhythms of the adult nematode using a bioluminescence-based circadian transcriptional reporter. We show that mutations of *lin-42* and *kin-20* generate a significantly longer endogenous period, suggesting a role for both genes in the nematode circadian clock, as in other organisms. These phenotypes can be partially rescued by overexpression of either gene under their native promoter. Both proteins are expressed in neurons and seam cells, a population of epidermal stem cells in *C. elegans* that undergo multiple divisions during development. Depletion of LIN-42 and KIN-20 specifically in neuronal cells after development was sufficient to lengthen the period of oscillating *sur-5* expression. Therefore, we conclude that LIN-42 and KIN-20 are critical regulators of the adult nematode circadian clock through neuronal cells.

## Introduction

Circadian rhythms are endogenous oscillations that allow adaptation to our cyclic environment, maintaining an intrinsic periodicity close to 24 hours. Maintained under constant conditions as free running rhythms (FR), circadian rhythms are synchronized by environmental signals (‘time givers’, or zeitgebers) such as light and temperature, and their period remains relatively unchanged by variations in temperature [1]. In metazoans, circadian rhythms are modulated by central and peripheral clocks that depend on the activity of a group of evolutionarily conserved clock genes that regulate interlocked transcription-translation feedback loops (TTFL) [2]. In mammals, CLOCK and BMAL1 proteins form a heterodimer that acts as a positive element in the core feedback loop activating the transcription of clock genes, such as PERIOD (PER) and CRYPTOCROME (CRY), by binding to the E-boxes in their promoters. In turn, PER binds to CRY and CK1*ε/δ*, forming multimeric complexes that function as the negative element in the feedback loop by inhibiting the activity of CLOCK:BMAL1, thereby hindering their own transcription and closing the TTFL [3–5]. PER proteins are progressively phosphorylated by CK1ε/δ to regulate its stability, resulting in daily changes in PER levels and CLOCK/BMAL1 activity [6].

Circadian rhythms of physiological and behavioral variables, such as gene expression and locomotor activity, have been observed in the nematode *C. elegans* [7]; however, the molecular mechanism of its central pacemaker is currently unknown. With the exception of *cry* (*cryptochrome*), *C. elegans* has homologs to other metazoan clock genes such as *lin-42* (*period*), *kin-20* (CK1ε/δ), *aha-1* (*cycle/bmal1*), *nhr-23* (*ror□/□*), *nhr-85* (*rev-erb□/β*), and *nhr-25* (*sf-1*) [8–11]. Some of these genes participate in other functions in the nematode, like the regulation of developmental timing [12–14], but their function in circadian rhythms has been understudied at the molecular level.

LIN-42 has seven different isoforms (isoforms a-g) that are relevant in the development of the nematode to different extents. For example, the absence of *lin-42a* (allele *ok2385*) or *lin-42b* (allele *ox461*) generates larval arrest and development defects [12]. By contrast, the absence of *lin-42c* (allele *n1089*) generates a less severe phenotype [15] and little is known about other isoforms. *lin-42* is expressed in epidermal cells, seam cells (multipotent lateral epidermal cells that undergo multiple divisions during larval development), the pharynx, and neuronal cells [12, 16]. In the complete absence of *lin-42*, hypodermal seam cells, vulval precursor cells, and the migration of sex myoblasts all develop precociously [17]. *lin-42* mRNA levels oscillate during each of the four molting cycles of postembryonic development (approximately 8-10 h at 25°C) [16] regulating developmental timing and entrance into an alternative dauer larval stage [12, 15, 17] likely through gene expression modulation [18–21]. The *kin-20* gene also encodes seven isoforms (isoforms a-g), and it is expressed in seam cells and neurons [8]. Dysfunctional KIN-20 proteins cause developmental and molting defects [8, 22]. KIN-20 also regulates the expression of *lin-42* in the larval stages [21]. In adult nematodes, KIN-20 stabilizes nervous system architecture after axon outgrowth [22].

Previous studies suggested that LIN-42 and KIN-20 are involved in nematode circadian rhythms. *lin-42* loss-of-function mutants show defects in circadian rhythms of locomotor activity [23], as is the case for PER mutants in other metazoa [24, 25]. Likewise, pharmacological inhibition of KIN-20 activity with the CKIδ/ε-selective kinase inhibitor PF-670462 significantly lengthens the period of *sur-5* bioluminescence rhythms in nematodes [26], similar to its effects on clocks in diverse species, including plants, *Drosophila* and mammals [27–29]. However, the functional roles of LIN-42 and KIN-20 in adult circadian rhythms have not been deeply characterized at the molecular level to date. This is due, in part, to the complexity of the different isoforms from both genes and the lack of tools to investigate the functional role of these proteins in adult circadian rhythms. We hypothesized that LIN-42 and KIN-20 play an important role in the regulation of circadian rhythms in *C. elegans*, similar to what occurs in other models such as mammals and *Drosophila*. We tackled this by using complementary approaches, combining biochemistry with genetic manipulations targeting the different isoforms of LIN-42 and KIN-20, and protein degradation post-development.

In this study, we characterized the effects of *lin-42* and *kin-20* disruption on circadian rhythms in adult nematodes by means of a sensitive bioluminescence-based reporter system [26]. We observed significant changes in the luminescent circadian rhythms driven by the *sur-5* promoter in *lin42(ox461)*, *kin-20(ok505)*, and *kin-20(ox463)* mutants. Joint neuronal depletion of both LIN-42 and KIN-20 proteins after development using an auxin-inducible degradation (AID) system altered the transcriptional rhythms of *sur-5*. This effect was absent, however, when the proteins were depleted in seam cells, highlighting the specificity of these genes in neurons to regulate circadian rhythms. We also show that the quaternary architecture of LIN-42 structure is conserved with mammalian PER2, suggesting that LIN-42 and KIN-20 may retain some functional similarity in regulation of the circadian clock of adult nematodes.

## Results

### The quaternary architecture of LIN-42 is structurally similar to mouse PER2

LIN-42 has long been established as a homolog to PER [16], which has essential roles in metazoan circadian rhythms. While *M. musculus* and *D. melanogaster* PER have a similar function as transcriptional co-repressors in the TTFL, the molecular details of the *Drosophila* TTFL differ from most other metazoans, which are genetically wired more similar to mammalian clocks [30]. Mammalian PER proteins are largely intrinsically disordered but contain four domains necessary for their circadian function: two tandem PAS domains (Period-Arnt-Sim, PAS-A and PAS-B) that homodimerize PER, a CK1 binding domain (CK1BD), and a CRY-binding domain (CBD). The longest isoform of LIN-42 is approximately half the length of mouse PER2 (598 vs. 1225 amino acids) and it has conserved PAS and CK1 binding domains [17] (Figure 2A).

The PER proteins from *Drosophila* and mouse exhibit different modes of dimerization by their PAS-B domains [31–33], which may influence their function at the molecular level. The LIN-42 PAS-B domain is 24.4% and 24.0% identical to mouse PER2 and *Drosophila* PER, respectively (Figure 1A). To gain insight into the function of the *C. elegans* PAS domains, we determined the structure of the LIN-42 dimer. Although we crystallized the protease-resistant core of the LIN-42 N-terminus (Figure S1A-B), density was poor for the putative N-terminal PAS domain (residues 41-146), which deviates from ideal PAS domain topology [34] (Figure S1C). However, we were able to build a model of the LIN-42 PAS-B dimer resolved to 2.4 Å (Figure 1A, Supplementary table 1) that closely resembles the mouse PER2 PAS-B dimer (Figure 1B). This was similar to the structural prediction by AlphaFold, which had two tandem PAS domains at high confidence followed by intrinsic disorder (Figure S1D-E). Altogether, this demonstrates a conserved dimerization strategy of LIN-42/PER among worms and mammals that may manifest in functional similarities as well.

**Figure 1.**
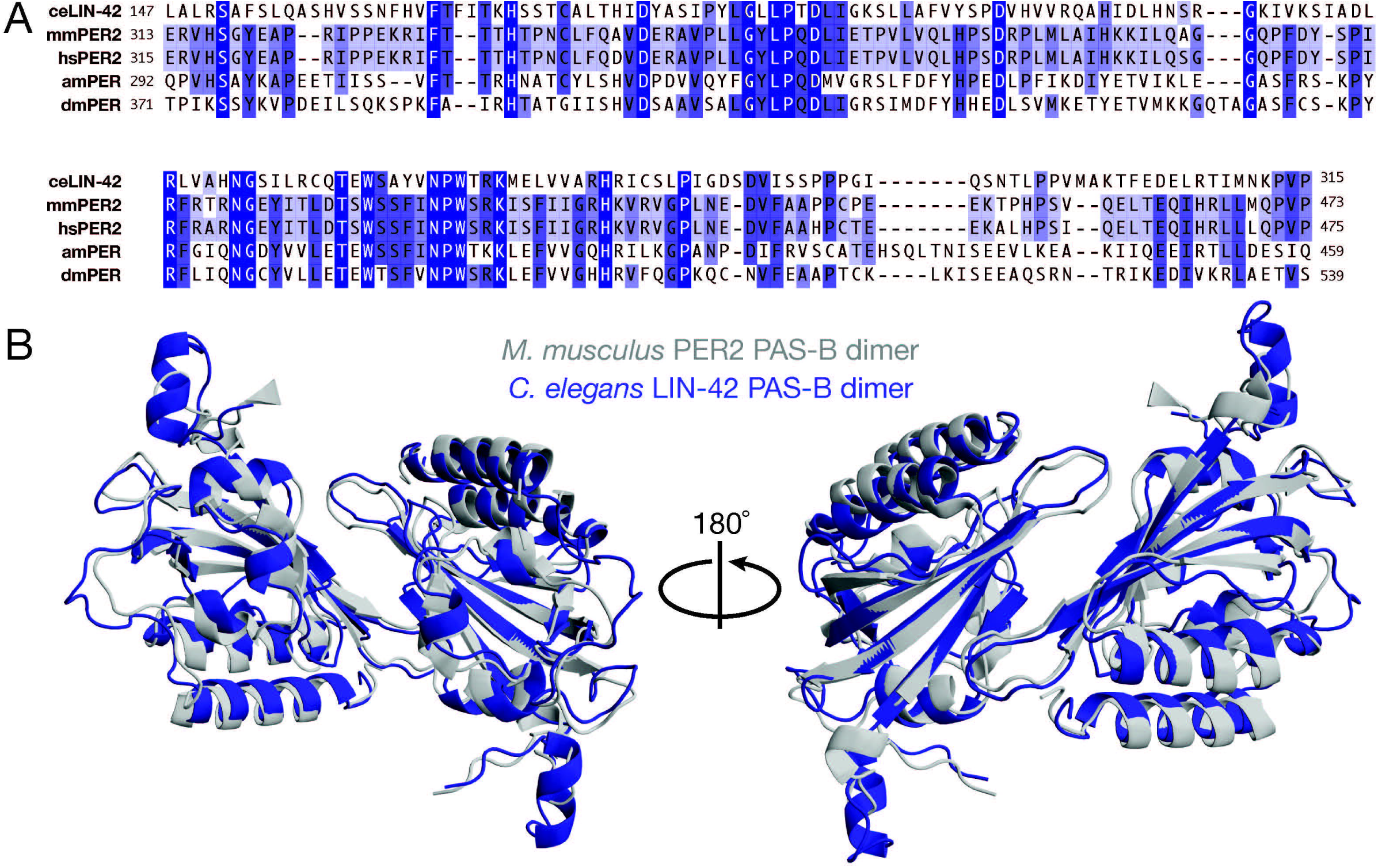
Conservation of mammalian PER quaternary structure in *C. elegans* LIN-42. **A.** Sequence alignment of *C. elegans, M. musculus, H. sapiens, A. mellifera,* and *D. melanogaster* PER PAS-B domains. **B.** Structural alignment of *C. elegans* LIN-42 PAS-B dimer (PDB 8GCI, blue) and *M. musculus* PAS-B dimer (PDB 3GDI, gray).

**Figure 2.**
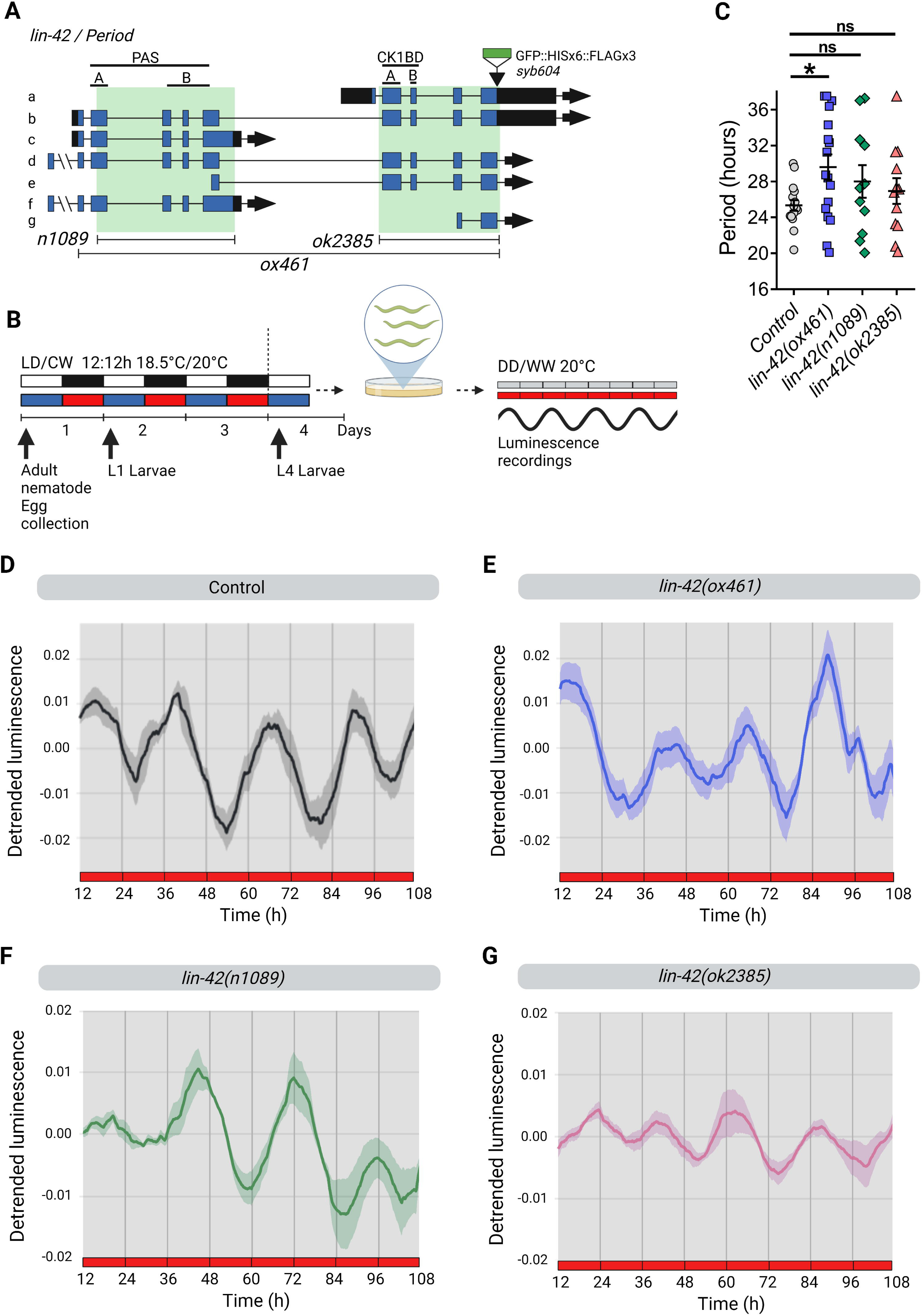
LIN-42 modulates period length under constant conditions. **A.** The *C. elegans lin-42* locus. The *n1089* allele is a deletion of the PAS-A and B domains, the *ok2385* allele is a mutation that affects the CK1BD (Casein Kinase 1 Binding Domain) and the *ox461* allele is a full deletion that lacks both (PAS and CK1BD). **B.** Schematic assay of the luminescence experiments and photo/thermoperiodic conditions. Black/white bars indicate dark/light; blue/red bars indicate cold/warm; gray and red bars indicate the FR conditions. Nematodes were grown under dual cyclic conditions of 12h:12h Light/Dark (LD, ∼150/0 µmol/m^2^/s) and Cold/Warm (CW, 18.5°C/20°C); ZT0, lights on and cold temperature phase onset, on NGM plates with a lawn of *E. coli* HB101. Embryos were collected from adult nematodes, then L1 (larval stage 1) nematodes were grown on NGM plates with bacteria for 3 days. Once at the L4 stage, approximately 50 nematodes were transferred to the liquid luminescence media. Luminescence assays were performed under FR conditions (DD/WW, 20°C) for 7 days. **C.** Average endogenous period was determined for hours 18h to 38h for nematodes exhibiting a rhythm in FR. Error bars represent SEM. The average endogenous period of mutant strains of *lin-42* was compared with the control strain (25.34 ± 0.60 h, n=16): *ox461* mutant (29.59 ± 1.41 h, n=17), *n1089* mutant (28.00 ± 1.81 h, n=11) and *ok2385* mutant (26.94 ± 1.43 h, n=12), *p=0.0446; ns, p>0.05, one-way ANOVA followed by Dunnett’s multiple comparisons test. The analysis includes three biological replicates for each strain. Average reporter activity of rhythmic adult populations under FR conditions (DD, 20°C), control (**D**) and *lin-42(ox461)* (**E**), *lin-42(n1089)* (**F**), *lin-42(ok2385)* (**G**). Luminescence signals are shown as mean ± SEM. The average reported activity was displayed with populations showing a similar first peak, representative single traces are shown in figure S2. Each population consisted of 50 adult nematodes per well.

### LIN-42 modulates period length and circadian entrainment in adult nematodes

To probe the functional role of LIN-42 in the *C. elegans* circadian system, we used the luciferase reporter *psur-5::luc::gfp* as a clock output [26], integrating the construct into the nematode genome by UV radiation (control strain *qvls*8, Table 1). We then introduced the reporter into *lin-42* mutant backgrounds by genetic crosses with this control strain. First, we tested three different mutants for LIN-42 (Figure 2A) that each delete various domains important for PER functions in mammals (Supplementary Table 2). *lin-42(ox461)* is a null allele with a 10226 bp deletion lacking the N-terminal PAS-A and B domains and the C-terminal CK1BD [5, 35–37]. The CKBD possesses two conserved motifs previously referred to as the SYQ and LT domains by Edelman TL et al. 2006 [17], now termed CK1BD-A and CK1BD-B [35]. We also tested the effects of *lin-42(n1089)* that has a deletion of 5233 pb, affecting only the PAS domains, and *lin-42(ok2385)* that has a 2632 pb deletion, missing the CK1BD domain and downstream C-terminus.

**Table 1.** PDB.

Data Collection and Refinement Statistics:
|  | Lin42 |
| --- | --- |
| **Data collection** |  |
| Beam line | APS (23IDD) |
| Space group | I 2 2 2 |
| Unit cell dimensions<br>(a,b,c) (Å), ($\alpha$, $\beta$, $\gamma$) (°) | 50.73, 105.9, 152.9, 90 90 90 |
| Resolution Range (Å)<br>(highest shell)* | 45.94 - 2.42 (2.51 - 2.42) |
| Wavelength (Å) | 1.033 |
| Total observations | 114830(11082) |
| Unique reflections | 16123 (1634) |
| Completeness (%) | 99.52 (98.48) |
| $R_{\text{merge}}$ | 14(65.8) |
| $\langle I/\sigma \rangle$ | 10.2(3.3) |
| CC1/2 | 99.4(89.5) |
| Redundancy (highest shell) | 7.1(6.8) |
| **Refinement** |  |
| $R_{\text{work}}$ %/ $R_{\text{free}}$ % | 26.5 (29.5) |
| Number of non-hydrogen atoms | 1394 |
| Protein Residues | 169 |
| Ligand/ion | - |
| Water | 76 |
| B-factor (Wilson) | 32.4 |
| RMSD Bond length (Å) | 0.008 |
| RMSD Bond angle | 1.25 |
| Ramachandran favored(%) / Ramachandran outliers (%) | 97.0/0.0 |

\* Values in parenthesis are for higher resolution shell.

**Table 2.**
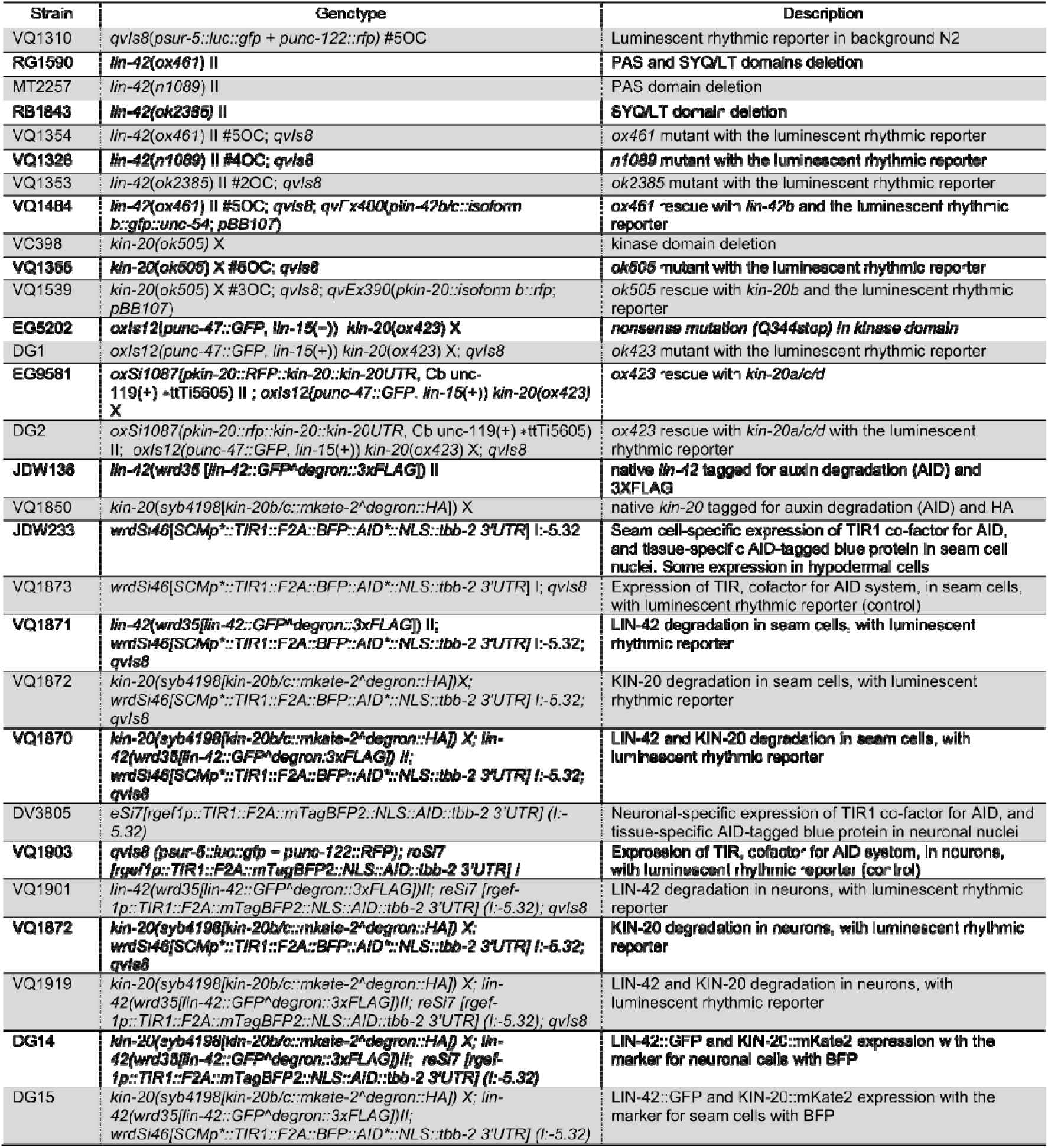
Strains.

**Table 3.**
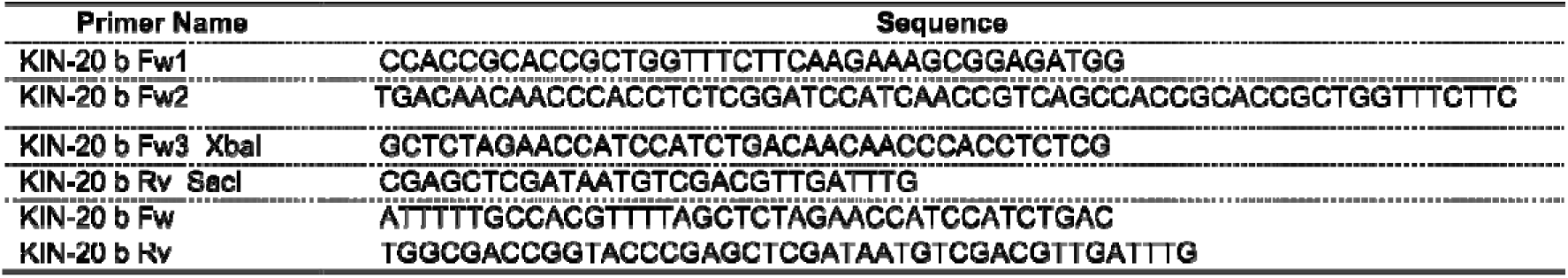
Primers.

Animals were synchronized to a dual LD/CW cycle (∼150/0 µmol/m2/s; 18.5°C/20°C) and the bioluminescent activity of the *sur-5* promoter was measured thereafter in adult nematodes in constant conditions of darkness and 20°C (DD/WW) as described in Goya et al. [26] (Figure 2B). Rhythmic *sur-5* promoter activity occurred in all strains under constant conditions, indicating that the nematode exhibits circadian rhythms under free running (FR) (Figure 2D-G and S2A-I). Although there was considerable higher variability of period across the mutant populations, we found that *lin-42(ox461)* animals showed a significantly longer period than the control strain (29.59 ± 1.41 h, n = 17 mutant vs 25.34 ± 0.60 h, n=16 control) (Figure 2C-E and S2B). However, no significant difference was observed in the mutants of the PAS domains (Figure 2C, 2F and S2C, 27.1 ± 1.74 h, n=10 mutant vs control) and the CK1BD domain (Figure 2C, 2G and S2D, 26.94 ± 1.43 h, n=12 mutant vs control). Given that an abnormal circadian phenotype manifests only when both PAS and CK1BD domains are absent, this suggests that the presence of either of them is sufficient for circadian rhythm regulation.

To further characterize the effect of the *lin-42* full deletion on clock synchronization, we analyzed luminescence rhythms first under cyclic and then constant conditions. We measured the luminescence of adult nematode populations over 3 days in LD/CW cycles (∼150/0 µmol/m^2^/s; 15.5°C/17°C) and then for 4 days under constant conditions (DD/WW, 17°C) [26] (Figure 3A). Both the control strain and the *lin-42(ox461)* strain exhibited a *sur-5* promoter rhythm under cyclical and FR conditions, thereby providing additional evidence that nematodes can be synchronized through a dual cycle of light and temperature (Figure 3C-D and S3B). Consistent with our previous experiments, the *lin-42(ox461)* mutant strain exhibited a longer period under constant conditions compared to the control strain (26.41 ± 0.47 h, n=32 mutants vs 24.4 ± 0.57 h, n=37 control) (Figure 3B). The *lin-42(ox461)* strain exhibited lower synchronization following this entrainment protocol compared to the control strain, as evidenced by acrophase shift of the Rayleigh in LD/CW and DD/WW conditions, suggesting that a masking mechanism is involved (i.e., a direct effect of the zeitgeber on circadian rhythms) (Figure 2F).

**Figure 3.**
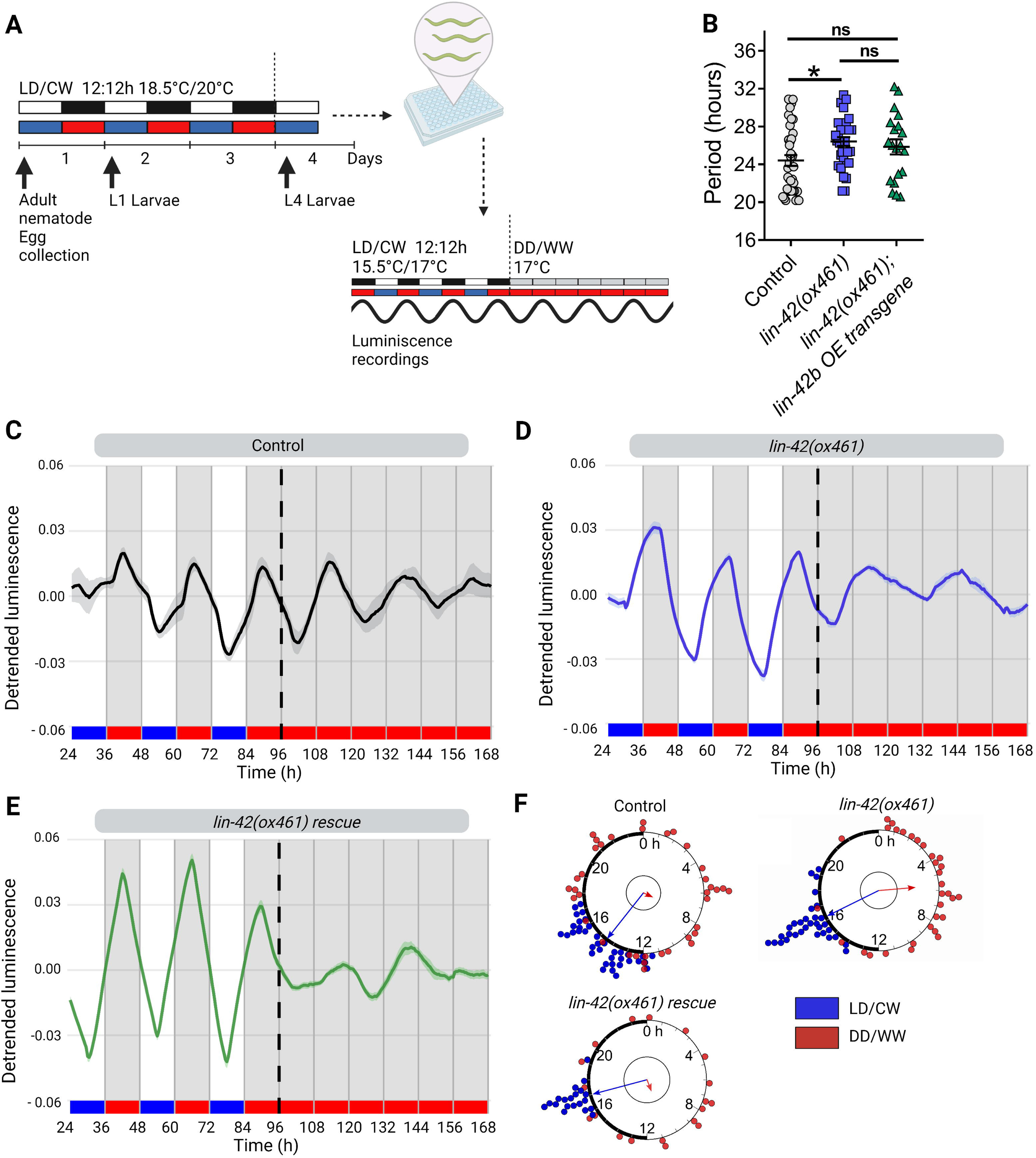
LIN-42 modulates circadian entrainment in adult nematodes. **A.** Schematic assay of the luminescence experiments and photo/thermoperiodic conditions. Black/white bars indicate dark/light; blue/red bars indicate cold/warm; grey and red bars indicate the FR conditions. Nematodes were grown under dual cyclic conditions of 12h:12h Light/Dark (LD, ∼150/0 µmol/m^2^/s) and Cold/Warm (CW, 18.5°C/20°C); ZT0, lights on and cold temperature phase onset, on NGM plates with a lawn of *E. coli* HB101. Embryos were collected from adult nematodes, then L1 (larval stage 1) nematodes were grown on NGM plates with bacteria for 3 days. Once at the L4 stage, approximately 50 nematodes were transferred to the liquid luminescence media. Luminescence assays were performed under dual cyclic conditions 12h: 12h of Light/Dark (LD, ∼150/0 µmol/m^2^/s) and Cold/Warm (CW, 15.5°C/17°C) for 3 days; ZT0, lights on and cold temperature phase onset. Then, the luminescence was measured for 4 more days under FR conditions (DD/WW, 17°C). **B**. Average endogenous period of rhythmic populations of control (24.4 ± 0.57 h, n=37), *ox461* mutants (26.41 ± 0.47 h, n=32) and *lin-42b* overexpression (OE) transgene strain (25.85 ± 0.79 h, n=20). One-way ANOVA followed by Turkey’s multiple comparisons test, *p=0.0302; ns, p>0.05. Representative luciferase activity rhythms of adult populations under dual cyclic conditions (LD/CW, ∼150/0 µmol/m^2^/s; 15.5°C/17°C) and FR conditions (DD, 17°C), control (**C**), *lin-42(ox461)* (**D**) and *lin-42b* transgene (**E**). Luminescence signals are shown as mean ± SEM. The average reported activity was displayed with rhythmic populations showing a similar first peak in FR. Each population consisted of 50 adult nematodes per well and the analysis includes three biological replicates for each strain. The reported activity from all the wells for each strain is displayed in Figure S3. **F.** Rayleigh plots showing the phase of the bioluminescent peak under dual cyclic conditions (LD/CW, blue dots) and the first bioluminescent peak on the first day of release to FR (DD/WW, red dots) for rhythmic population nematodes: Control (LD/CW: 14.55 ± 0.27 h, n=37; R=0.90 and DD/WW: 7.85 ± 0.85 h, n=37; R=0.07), *lin-42(ox461)* mutants (LD/CW: 16.31 ± 0.20 h, n=32; R=0.95 and DD/WW: 5.68 ± 0.66 h, n=32; R=0.51) and *lin-42(ox461);lin-42b* OE transgene rescue (LD/CW: 17.03 ± 0.15 h, n=20; R=0.98 and DD/WW: 10.75 ± 1.09 h, n=20; R= 0.17). Arrows represent the average peak phase of *sur-5::luc* expression (mean vectors for the circular distributions) of each group. The length of the vector represents the strength of the phase clustering while the angle of the vector represents the mean phase. Individual data points are plotted outside the circle. The central circle represents the threshold for p=0.05.

We then tested whether a transgene of the longest *lin-42b* isoform, which encodes both PAS and CK1BD domains could rescue aberrant behavior in the *lin-42(ox461)* null mutant when overexpressed from an extrachromosomal array under its native promoter. Although this *lin-42b* transgene was not able to restore the period length to control values (Figure 3B and 3E), it improved entrainment to the wild-type level (Figure 3F). This partial rescue phenotype may be due to the nature of the extrachromosomal arrays, which induce high interindividual variability in gene expression. For *lin-42(n1089)* mutants that lack the PAS domain and *lin-42(ok2385)* mutants that lack the CK1BD domain, we did not find any significant differences in either period or synchronization compared to the control strain (Figure S4). Taken together, these results show that *lin-42* modulates the period of *sur-5-* driven luminescent rhythms and that either the PAS or the CK1BD domains may be sufficient for circadian entrainment in *C. elegans*.

### KIN-20 regulates circadian period in adult nematodes

To study the function of *kin-20* in the circadian system of *C. elegans*, we followed a similar approach as for *lin-42,* recording luminescence rhythms from two loss-of-function *kin-20* mutants (Figure 4A), both of which are dumpy, have egg-laying defects, and show progressive paralysis [22]. We examined luminescence rhythms under constant conditions (DD/WW, dark and 20°C) or under cyclic conditions (LD/CW, ∼150/0 µmol/m2/s; 15.5°C/17°C) followed by constant dark and temperature conditions (DD/WW, dark and 17°C). We first tested the null allele *ok505* that has a complete deletion of the kinase domain (2201 pb deletion, lacking exons 3-5) and an out-of-frame insertion (Figure 4A). Under constant conditions, *kin-20(ok505)* mutant animals showed a significantly longer period compared to the control strain (Figure 4B-D and S5A-B), 29.36 ± 1.05 h, n=26 mutant vs 24.31 ± 0.92 h, n=12 control). The *kin-20(ox423)* null mutant has a nonsense mutation (Q344stop) in the kinase domain that affects all isoforms (Figure 4A). Similar to *ok505, ox423* mutants also showed a significantly longer period in luminescent rhythms compared to control worms (Figure 4B-E and S5C, 30.22 h ± 1.47 h, n=11 mutant vs 24.31 ± 0.92 h, n=12 control).

**Figure 4.**
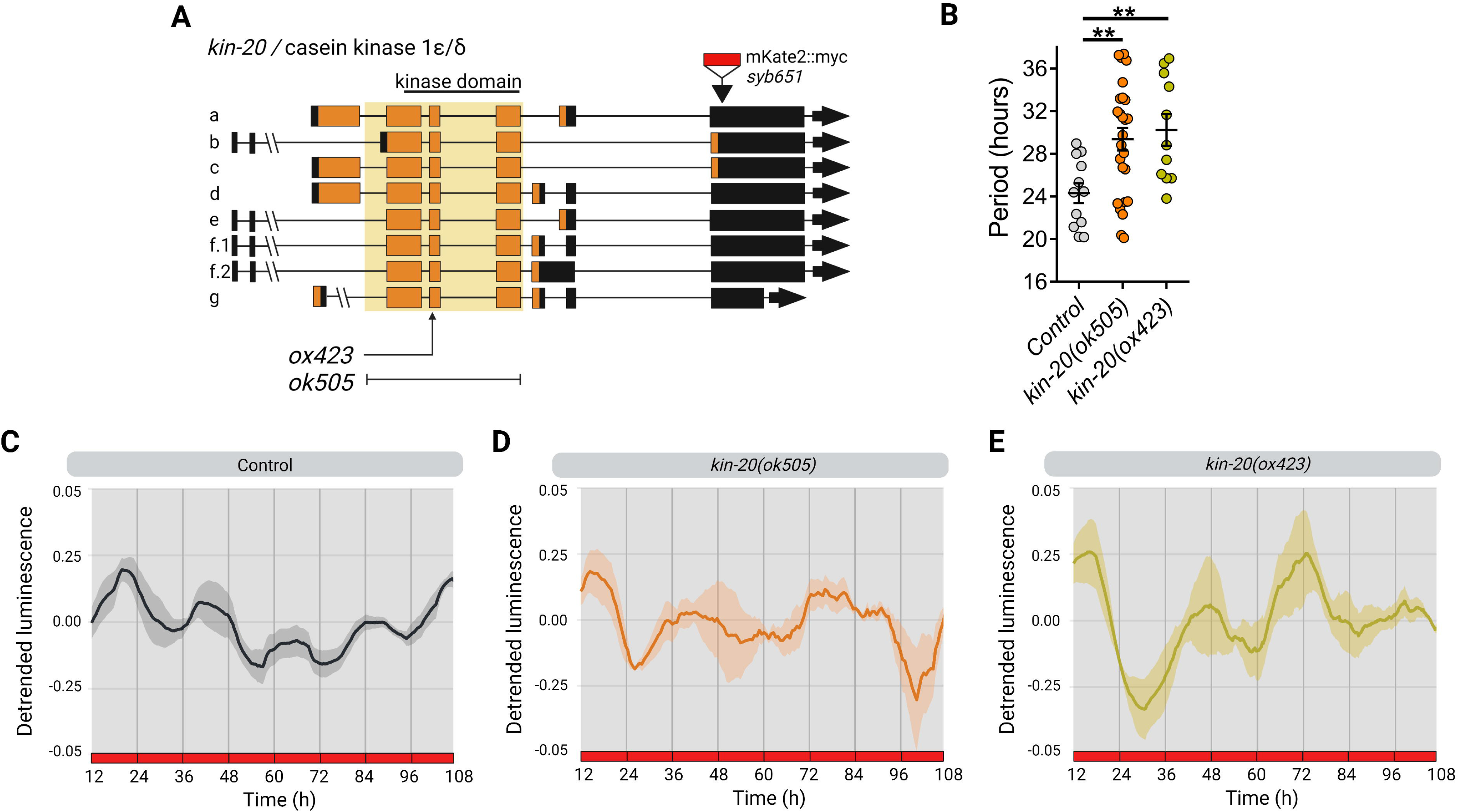
KIN-20 modulates period length in adult nematodes under FR conditions. **A**. The *C. elegans kin-20* locus. The *ok505* allele harbors a deletion mutation in the kinase domain and the *ox423* allele is a nonsense mutation (Q344stop) in the kinase domain. The *syb651* allele is a knock-in of *mKate2::myc* in the 3’ UTR of isoform b. **B.** Average endogenous period was determined for hours 18h to 38h for nematodes exhibiting a rhythm in FR. Error bars represent SEM. The average endogenous period of mutant strains of *kin-20* was compared with the control strain (24.31 ± 0.92 h, n=12): *ok505* mutants (29.36 ± 1.05 h, n=26) and *ox423* mutants (30.22 h ± 1.47 h, n=11), **p<0.01, one-way ANOVA followed by Dunnett’s multiple comparisons test. The analysis includes three biological replicates for each strain. **C**-**E.** Average reporter activity of rhythmic adult populations under FR conditions (DD, 20°C), control (**C**), *kin-20(ok505)* (**D**) and *kin-20(ox423)* (**E**). Luminescence signals are shown as mean ± SEM. The average reported activity was displayed with populations showing a similar first peak, representative single traces are shown in figure S5. Each population consisted of 50 adult nematodes per well.

We further analyzed the effect of *kin-20* mutants on luminescence rhythms of the nematode under cyclical conditions and release into free running conditions. The *kin-20(ok505)* strain exhibited rhythmic *sur-5* promoter activity under both cyclical and in FR conditions, similar to the control strain. Again, *kin-20(ok505)* animals showed a longer period compared to the control strain in FR (Figure 5A, E and S6A, 26.61 ± 0.69 h, n=39 mutant vs 24.4 ± 0.57 h, n=37 control). Next, we asked if overexpression of the isoform b of KIN-20 (Figure 4A), which has the highest homology to the kinase domain of CK1 ε/δ from mammals [9], was able to rescue the longer period phenotype of *ok505* mutants. For this, we generated a strain with an extrachromosomal array overexpressing *kin-20b* under its native promoter. This resulted in a partial rescue of the period compared to wild-type animals (Figure 5A, E and S6A, 25.73 ± 0.91 h, n=18 rescue vs 24.4 ± 0.57 h, n=37 control). All strains exhibited a similar dispersion of acrophases, showing low acrophase dispersion under cyclical conditions and higher acrophase dispersion in FR (Figure 5C). Consistent with the phenotypes for *kin-20(ok505)*, *kin-20(ox423)* mutants also showed a significantly longer period compared to the control (Figure 5B, F and S6B, 27.81 ± 0.92 h, n=18 mutant vs 24.93 ± 0.59 h, n=32 control). We then attempted to rescue *kin-20(ox423)* behavioral defects by generating animals carrying a single-copy of the transgene *oxSi1087* expressing RFP-tagged KIN-20, which includes all exons and 8.1 kb upstream of the first ATG codon (strain EG9581) [22]. These animals exhibited luminescence rhythms with a period length similar to wild-type controls (Figure 5B, F and S6B, 25.46 ± 0.89 h, n=24 rescue vs 24.93 ± 0.59 h, n=32 control). In addition, the ability to synchronize with the zeitgeber was comparable for all strains, as they displayed a similar dispersion of acrophases in LD/CW and DD/WW conditions (Figure 5D). Therefore, our data demonstrate that *kin-20* is necessary for determining the period of circadian rhythms in *C. elegans,* but is not involved in circadian entrainment.

**Figure 5.**
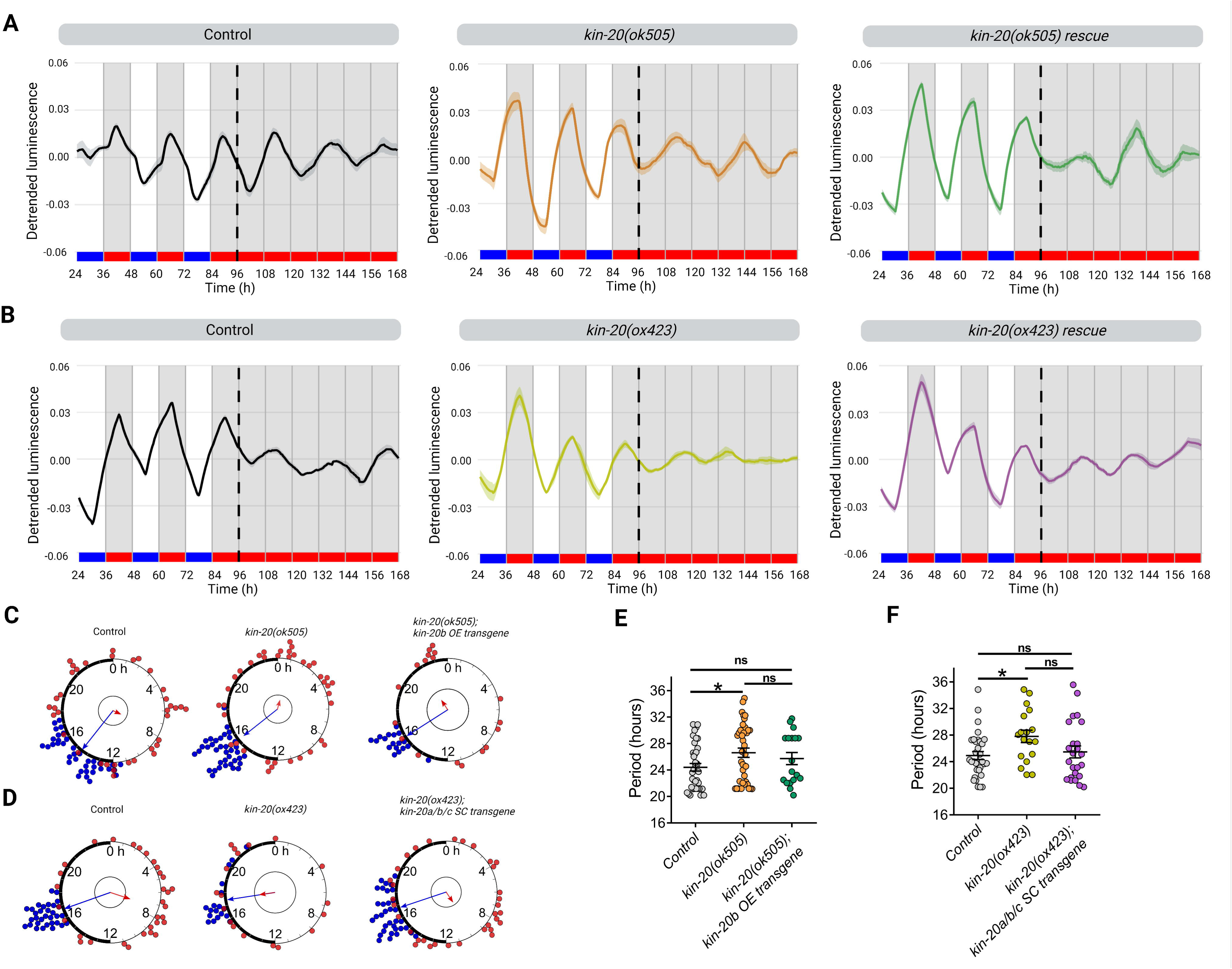
KIN-20 modulates period length in adult nematodes under cyclic conditions. **A-B.** Representative luciferase activity rhythms of adult populations under dual cyclic conditions (LD/CW, ∼150/0 µmol/m^2^/s; 15.5°C/17°C) and FR conditions (DD, 17°C): control, *kin-20(ok505)*, *kin-20b* overexpression (OE) transgene (**A**) and control, *kin-20(ox423)* and *kin-20a/b/c* single-copy transgene (**B**). Luminescence signals are shown as mean ± SEM. The average reported activity was displayed with rhythmic populations showing a similar first peak in FR. Each population consisted of 50 adult nematodes per well. The reported activity from all the wells for each strain is displayed in Figure S6. **C**-**D.** Rayleigh plots showing the phase of the bioluminescent peak under dual cyclic conditions (LD/CW, blue dots) and the first bioluminescent peak on the first day of release to FR (DD/WW, red dots) for rhythmic population nematodes: control (LD/CW: 14.55 ± 0.27 h, n=37; R=0.90 and DD/WW: 7.85 ± 0.85 h, n=37; R=0.07), *kin-20(ok505)* (LD/CW: 15.49 ± 0.12 h, n=39; R=0.97 and DD/WW: 22.83 ± 0.76 h, n=39; R=0.21), *kin-20b* overexpression (OE) transgene (LD/CW: 15.83 ± 0.18 h, n=18; R=0.97 and DD/WW: 2.03 ± 1.2 h, n=18; R=0.098) (**C**) and control (LD/CW: 16.7 ± 0.10 h, n=32; R=0.98 and DD/WW: 7.1 ± 0.76 h, n=32; R=0.35), *kin-20(ox423)* (LD/CW: 17.46 ± 0.37 h, n=18; R=0.91 and DD/WW: 17.43 ± 1.07 h, n=18; R=0.28) and *kin-20a/b/c* single-copy transgene (LD/CW: 16.08 ± 0.19, n=24; R=0.96 and DD/WW: 12.63 ± 0.94 h, n=24; R=0.25) (**D**). Arrows represent the average peak phase of *sur-5::luc* expression (mean vectors for the circular distributions) of each group. The length of the vector represents the strength of the phase clustering while the angle of the vector represents the mean phase. Individual data points are plotted outside the circle. The central circle represents the threshold for p=0.05. **E.** Average endogenous period of control (24.4 ± 0.57 h, n=37), *ok505* mutants (26.61 ± 0.69 h, n=39) and *kin-20b* overexpression (OE) transgene strain (25.73 ± 0.91 h, n=18). One-way ANOVA followed by Tukeyśs multiple comparisons test, *p=0.0421; ns, p>0.05. **F.** Average endogenous period of control (24.93 ± 0.59 h, n=32), *ox423* mutants (27.81 ± 0.92 h, n=18) and *kin-20a/b/c* single-copy (CS) transgene (25.46 ± 0.89 h, n=24). One-way ANOVA followed by Tukeyśs multiple comparisons test, *p=0.0365; ns, p>0.05.

### LIN-42b and KIN-20b regulate circadian rhythms in adult neurons

Although we have shown that both *lin-42* and *kin-20* are involved in circadian rhythm regulation in *C. elegans*, we cannot rule out the possibility of indirect developmental effects in the mutants. To address this and gain more insight into the tissues from which both genes act to modulate period length, we used the auxin-inducible degradation (AID) system [38]. Using CRISPR, we generated *lin-42b/c*::GFP::AID*::3xFLAG and *kin-20b/c*::mKate2::AID knock-ins. We crossed both of these AID knock-ins into a strain expressing the *TIR1* transgene, either in seam cells or in neurons, to drive the depletion of the targeted proteins (LIN-42 and KIN-20) in either cell type. These transgenes use a ribosomal 2A skip sequence to produce a nuclear localized BFP::AID* reporter, which enables measurement of both transgene expression and TIR1 activity (Figure 6A) [39]. We first analyzed the endogenous expression pattern of LIN-42B and KIN-20B in neurons that were labeled by *rgef-1p*::TIR-1::F2A::BFP::AID*::NLS::tbb-2 in L4 stage nematodes using confocal fluorescence microscopy. We found strong co-expression of LIN-42B and KIN-20B in neurons (Figure 6B, S7A). To further support this, we used the CeNGEN single-cell RNA sequencing dataset [40], which provides gene expression profiles of all 302 neurons of the *C. elegans* nervous system in L4 nematodes [41]. We found that the genes *lin-42* and *kin-20* are both expressed in pharyngeal neurons, motor neurons, sensory neurons, and interneurons (Figure S7C). Moreover, we generated a strain simultaneously carrying the two engineered knock-in loci, *lin-42b*::GFP::AID and *kin-20b*::mkate-2::AID, as well as the transgene *SCMp*::TIR-1::F2A::BFP::AID*::NLS::tbb-2, which expresses BFP in epidermal seam cells (Figure 6A). We found a strong co-expression of the proteins LIN-42B and KIN-20B in seam cells in L3/L4 stage larva as well (Figure 6C, S7B).

**Figure 6.**
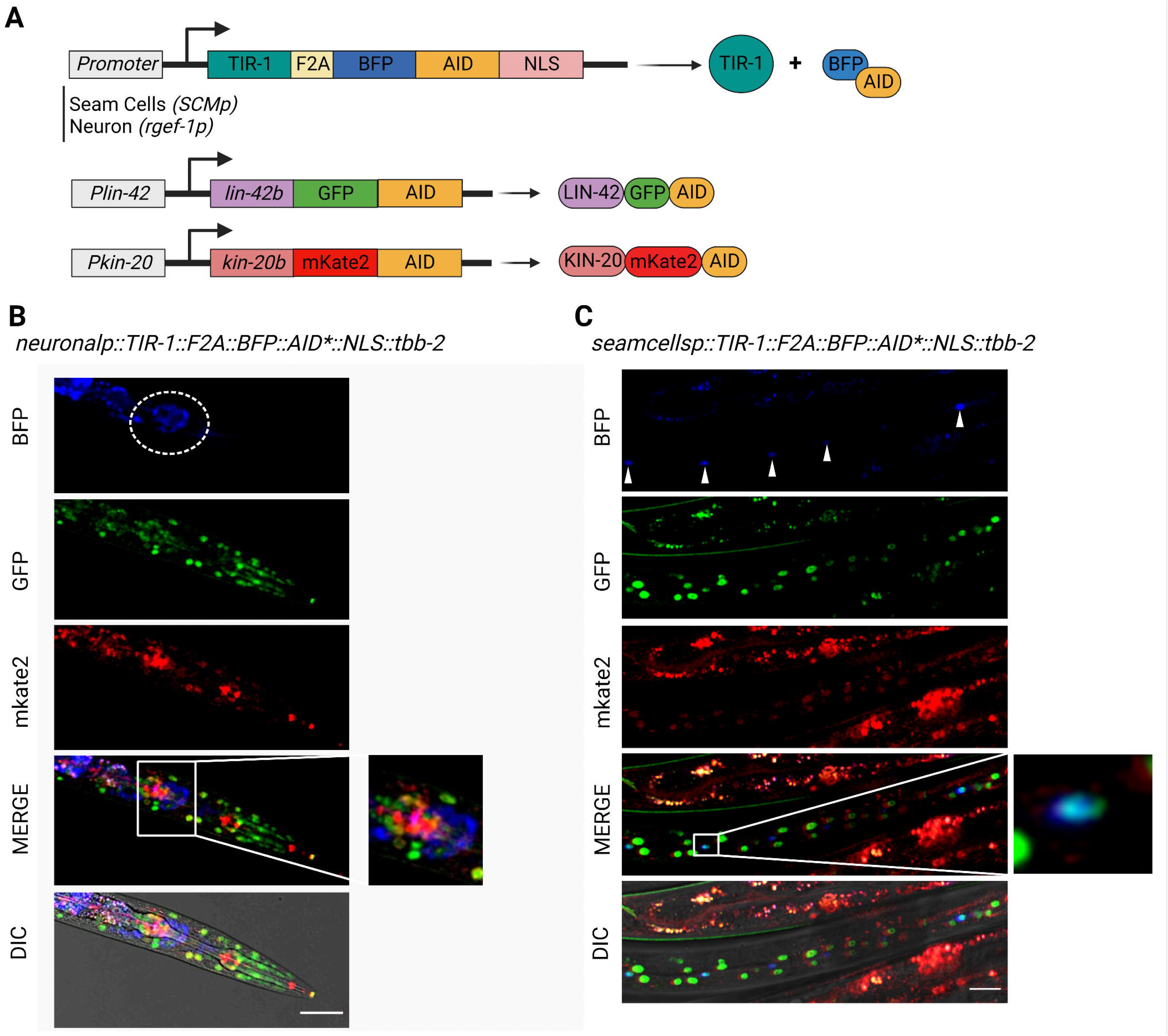
Expression of *lin-42b* and *kin-20b* in neurons and seam cells. **A**. Schematic of the auxin-inducible degron (AID) system for *lin-42b* and *kin-20b*. TIR-1 is expressed under the promoter *SCMp* in seam cells and *rgef-1* in neurons. The TIR-1::F2A::BFP::AID*::NLS transgene cassette encodes for two separate protein products: TIR-1, which will interact with endogenous SCF proteins to produce an E3 ubiquitin ligase complex and can only bind the AID sequence in the presence of auxin; and an AID*-tagged BFP protein with a c-Myc nuclear localization signal (NLS) that functions as a readout for TIR-1 expression and internal control for TIR-1 activity. The sequence *lin-42b*::AID was tagged with GFP, and the sequence *kin-20b*::AID was tagged with mKate2. **B**. GFP and mKate2 are detected in the neurons BFP positive (white circle) in the L4 stage. **C**. GFP and mKate2 are detected in the seam cells BFP positive (arrowheads) in L3/L4 nematodes. BFP positives in gut granules is autofluorescence. Scale bars represent 20 µm.

Subsequently, we asked whether LIN-42B and/or KIN-20B act in the nervous system or seam cells to regulate circadian *sur-5p::luc* expression. First, we verified the degradation activity of TIR-1 in the tissues of interest under our luminescence recording conditions by following the fluorescence of BFP fused to TIR-1 (Figure 6A, S8A-B). For neurons, L4 stage worms were used to detect BFP-positive cells (Figure S8A). As seam cells are difficult to observe in adult nematodes, we used L3/L4 stage worms instead (Figure S8B). As expected, we observed a significant loss of BFP fluorescence in neurons or seam cells after 7 days of exposure to 4 mM auxin (K-NAA) [42], confirming tissue-specific degradation by TIR-1 (Figure S8). Together, these results demonstrate that TIR-1 can efficiently deplete AID-tagged proteins in neuronal or seam cells under our luminescent recording conditions.

Next, we analyzed whether the fusion of GFP::AID or mKate2::AID alters the function of LIN-42B or KIN-20B, respectively. We measured transcription of the reporter gene *sur-5::luc::gfp* in animals with the *lin-42b*::GFP::AID and *kin-20b*::mKate2::AID constructs (with TIR-1 expressed in neurons or seam cells) and compared them with the control strain (*qvIs8*). We analyzed luminescence rates from L4 stage transgenic nematodes for 3 days under cyclic conditions (LD/CW, 15.5/17°C, 12:12 h) and then for 4 more days under constant conditions (DD/WW, 17°C). The transgenic nematodes expressing *lin-42b*::GFP::AID and *kin-20b*::mkate2::AID, crossed with either TIR-1 expressed in neuronal cells (Figure S9A, B) or seam cells (Figure S9C, D), showed similar behavior to the control strain in the absence of auxin. Thus, these results indicate that the modification of the *lin-42* and *kin-20* alleles by the fusion of the fluorescent protein and the AID sequence and the expression of TIR-1 do not alter nematode rhythms.

To examine the effect of depleting KIN-20 and LIN-42 in either seam cells or nervous system on the expression of *sur-5::luc::gfp*, we measured luminescence rates in adult transgenic nematodes for 3 days under cyclical conditions (LD/CW, 15.5/17°C, 12:12 h) and then for 4 more days under constant conditions (DD/WW, 17°C). We added 4 mM K-NAA in the luminescence medium on the first day of recording and we compared the endogenous periods in transgenic nematodes exposed to drug or vehicle. Depletion of LIN-42 in neuronal cells, although showing a trend towards an altered period, did not generate a significant change in the period (Figure 7A, B, 26.32 ± 0.89 h, n=19 LIN-42::AID+4 mM K-NAA vs 24 h ± 0.86, n=11 control+4 mM K-NAA). However, we did observe a significant lengthening of period in nematodes with depletion of KIN-20 in neurons (Figure 7A, C, 28.32 ± 1.12 h, n=21 KIN-20::AID+4 mM K-NAA vs 24 ± 0.86 h, n=11 control+4 mM K-NAA). These auxin-dependent circadian phenotypes in nematodes expressing TIR-1 in neuronal cells were similar to the phenotype of *kin-20(ok505)* and *kin-20(ox423)* mutants (Figure 5). A depletion of both proteins (KIN-20 and LIN-42) in neurons also generated a significant change in period (Figure 7A, D, 30.01 ± 0.70 h, n=12 LIN-42/KIN-20::AID+4 mM K-NAA vs 24 ± 0.86 h, n=11 control+4 mM K-NAA). The longest circadian period was observed when both KIN-20 and LIN-42 proteins were depleted, compared to nematodes where only KIN-20 or LIN-42 was depleted. Noteworthy, we did not observe any difference in period in worms depleted of LIN-42B and KIN-20B in seam cells, suggesting that the transcription of circadian *sur-5* is not regulated by KIN-20 or LIN-42 proteins in this of cell type (Figures 7E-H). Collectively, these data show that KIN-20 and LIN-42 regulate the period of *sur-5* luminescent rhythms in the nervous system of adult worms, consistent with post-transcriptional or post-translational regulation of a clock located in *C. elegans* neurons, as in other organisms [4, 43].

**Figure 7.**
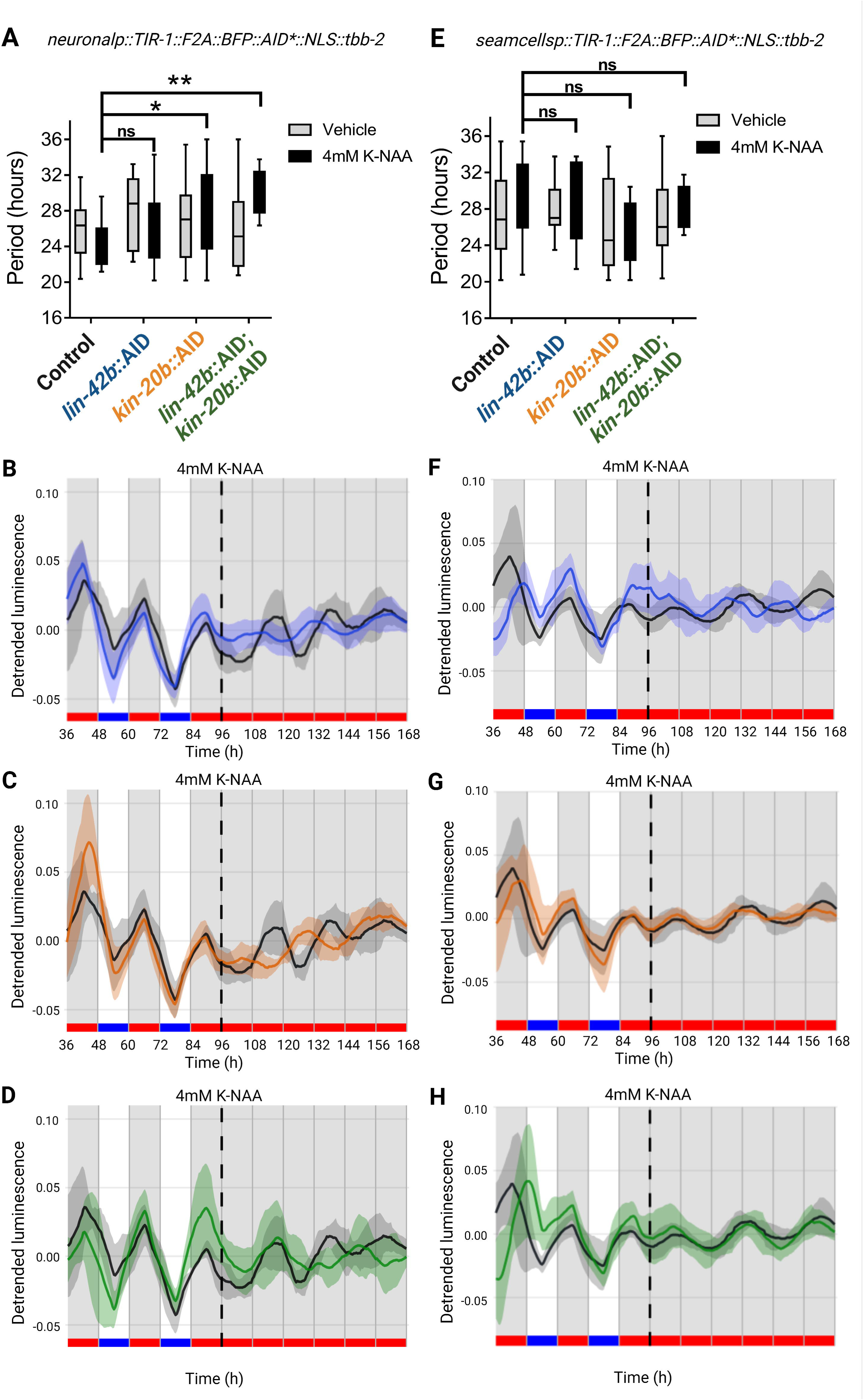
LIN-42b and KIN-20b regulate circadian rhythms in adult neurons. **A**. Average endogenous period of adults nematodes that express *rgef-1p*::TIR-1::F2A::BFP::AID*::NLS::tbb-2 and the respective AID-tagged protein, exposed vehicle or 4 mM Auxin continuously throughout the luminescence experiment. Nematodes exposed to vehicle: Control (25.97 ± 0.85 h, n=17), *lin-42b*::AID mutants (27.62 ± 1.14 h, n=13), *kin-20b*::AID mutants (26.93 ± 1.30 h, n=12) and *lin-42b*::AID; *kin-20b*::AID mutants (26.07 ± 0.92, n=25). Nematodes exposed to 4 mM Auxin: Control (23.99 ± 0.86 h, n=11), *lin-42b*::AID mutants (26.31 ± 0.89 h, n=19), *kin-20b*::AID mutants (28.32 ± 1.12 h, n=21) and *lin-42b*::AID; *kin-20b*::AID mutants (30.00 ± 0.70 h, n=12). Two-way ANOVA followed by Dunnett’s multiple comparisons test, *p=0.0160, **p=0.0020; ns, p>0.05. **B**-**D**. Representative luciferase activity rhythms of adult populations shown in E, exposed to 4mM Auxin (K-NAA), under the same dual cyclic conditions and FR conditions. Luminescence signals are shown as mean ± SEM. The average reported activity was displayed with rhythmic populations showing a similar first peak in FR. Each population consisted of 50 adult nematodes per well and the analysis includes three biological replicates for each strain. **E**. Average endogenous period of adults nematodes that express *SMCp*::TIR-1::F2A::BFP::AID*::NLS::tbb-2 and the respective AID-tagged protein, exposed to vehicle or 4 mM Auxin continuously throughout the luminescence experiment. Nematodes exposed to vehicle: Control (26.96 ± 1.05 h, n=18), *lin-42b*::AID mutants (27.87 ± 0.74 h, n=14), *kin-20b*::AID mutants (26.16 ± 1.12 h, n=17) and *lin-42b*::AID; *kin-20b*::AID mutants (26.88 ± 1.11, n=16). Nematodes exposed to 4 mM Auxin: Control (28.87 ± 1.02 h, n=18), *lin-42b*::AID mutants (28.86 ± 1.60 h, n=8), *kin-20b*::AID mutants (25.44 ± 1.04 h, n=10) and *lin-42b*::AID; *kin-20b*::AID mutants (27.93 ± 0.71, n=11). Two-way ANOVA followed by Sidak’s multiple comparisons test, ns, p>0.05. **F**-**H**. Representative luciferase activity rhythms of adult populations shown in **E**, exposed to 4mM Auxin (K-NAA), under the same dual cyclic conditions and FR conditions. Luminescence signals are shown as mean ± SEM. The average reported activity was displayed with rhythmic populations showing a similar first peak in FR. Each population consisted of 50 adult nematodes per well and the analysis includes three biological replicates for each strain. The reported activity from all the wells for each strain is displayed in Figure S10.

## Discussion

PER and CK1ε/δ proteins participate in the regulation of circadian rhythms throughout metazoans [5, 25] and have homologous genes in *C. elegans*, LIN-42 and KIN-20 respectively. We examined whether these proteins are also involved in regulating *C. elegans* circadian rhythms. We measured bioluminescence rhythms of the *sur-5* reporter gene, which allowed for continuous, non-invasive recording in the nematode [26]. Circadian regulation of the gene *sur-5* can be synchronized by a dual cycle of light and temperature and is maintained under constant conditions of light and temperature with a period close to 24 h, while also exhibiting temperature compensation [26], the three defining features of circadian rhythms.

A disruption in PER generates alterations in the period or eliminates rhythms altogether in mice and flies [24, 44–46]. Consistent with this, we found that the *lin-42(ox461)* null mutant lengthens the period of transcriptional rhythms at the *sur-5* promoter and produces a decrease in the percentage of nematodes that can be entrained by external stimuli, such as temperature and light. However, mutations exclusively in the PAS domains (*lin-42 (n1089)*) or the CK1BD domain (*lin-42 (ok2385)*) did not generate significant alterations in the endogenous period. Our observation that the *lin-42b* transgene did not rescue the long period of the null allele could be due to a mosaic expression of the transgene, which normally occurs in rescue assays of *C. elegans* mutants with extrachromosomal arrays [47]. In mice, *Per1* overexpression under a constitutive promoter lengthens the period of activity and body temperature rhythms [48], suggesting that LIN-42 may have a distinct mechanism in timekeeping from its mammalian homologs.

*lin-42* isoforms are characterized by the inclusion of different domains: *lin-42a* contains the CK1BD domain, *lin-42c* contains the PAS-A and PAS-B domains, and *lin-42b* has both [17]. During development, *lin-42b* and *lin-42a* play a more important role than *lin-42c* [17, 49]. We found that isoforms containing both PAS and CK1BD domains, like *lin-42b/d*, seem to have a greater relevance in the circadian system of the adult nematode than isoforms containing only the PAS or the CK1BD domains, such as *lin-42c* or *a*, since specific deletions of these domains did not alter the endogenous period. This suggests that the different isoforms might regulate each other to generate the expression and proper function of *lin-42*; only deleting multiple domains induced a clear alteration in circadian rhythms. This is different than in mammals, where an in-frame disruption of the PAS domain alone in *mPer2^Brdm1^* mutant mice is enough to induce a significantly shorter period of locomotor activity and a loss of circadian rhythmicity in free running [36]. The *lin42b/c/a* isoforms exhibit transcriptional oscillations during developmental molt changes in *C. elegans* with ∼8 h periods at 25°C that are not temperature compensated [16]. In adult worms, previous studies did not find circadian transcriptional regulation of *lin*-*42* at the gene level under light/dark (LD) or cold/warm (CW) temperature cycles in whole-worm extracts [50, 51]. It would be interesting to see whether *lin-42* mRNA also oscillates during adulthood with a circadian pattern in a specific group of cells, in long-read RNAseq experiments to distinguish between different isoforms.

CK1ε/δ is important in maintaining the rhythmicity and period of the clock in mammals and other organisms [52, 53]. Here we show that the *kin-20(ok505)* and *kin-20(ox423)* mutants exhibit a significant lengthening of period compared to the control strain without affecting entrainment to external cues. KIN-20 function can be partially rescued by overexpression of *kin-20b* in the null *kin-20(ok505)* mutant or by single-copy expression of the isoforms a/b/c in *kin-20(ox423)*. This suggests that the function of KIN-20 is not only important for nematode development [8, 21] and neuronal stabilization [22]; in addition, it is relevant for maintaining rhythms in adult nematodes.

In mammals, CK1ε/δ stably binds to PER and phosphorylates it to regulate its stability via the proteasome [6, 37, 54]. While this particular interaction has not yet been demonstrated in *C. elegans*, it was recently shown that KIN-20 regulates the expression of LIN-42A in larval stages L3 and L4 [21]. Here, we show that LIN-42b and KIN-20b are co-expressed both in neuronal cells and in seam cells, although more research is needed to determine the possible role of other nematode casein kinases in the regulation of LIN-42 and a possible interaction between the proteins.

To study where is KIN-20 and LIN-42 activity necessary, we used the AID system to deplete these proteins in neurons or seam cells. Remarkably, we found a change in the period of *sur-5* transcription when LIN-42 and KIN-20 were depleted in neuronal cells, but not in epidermal seam cells. We observed the same when depleting just KIN-20 in neurons, but not when depleting just LIN-42 in the same group of cells. The exacerbated period change when depleting both proteins may be because the interaction between LIN-42 and KIN-20 is functionally different from mammals or that KIN-20 could regulate other proteins in nematodes that also participate in circadian rhythms.

In summary, we show that the PERIOD and CK1ε/δ clock homologs LIN-42 and KIN-20, play an important role in the generation of circadian rhythms in adult nematodes, mainly by regulating period length. Their evolutionarily and structural conservation of key domains as well as dimerization strategy for LIN-42 suggests that there may be some shared aspects of nematode and mammalian clocks. In addition, our results highlight the existence of an ancestral circadian clock in metazoans that relates the regulation of development and adult behavior in the nematode, strengthening its use as an exciting model system in chronobiology to explore the molecular origins of metazoan timekeeping.

## Materials and Methods

### Strains and maintenance

*C. elegans* strains were cultured as previously described [26]. All strains were maintained under a dual cycle of light and temperature, Light:Dark (LD, ∼150/0 µmol/m^2^/s) and Cold:Warm (CW, 18.5/20°C) cycles 12:12h, LD and CW were established in the best phase combination, as already described [26]. *C. elegans* hermaphrodites were analyzed.

Zeitgeber (i.e., “time giver” or entraining agent) time 0 or ZT0 (9:00 am) indicates the time of lights on and the cold phase. Circadian Time (CT) refers to a specific time in the free running conditions (constant darkness, DD, and warm constant temperature, WW). Photo and thermal conditions were controlled with an I-291PF incubator (INGELAB, Argentina) and temperature was monitored using DS1921H-F5 iButton Thermochrons (Maxim Integrated, USA). Four cool white LED light strips (approx.150 µmol/m^2^/s) were used as the light source for the luminescence recordings.

The mutant and transgenic strains used in this study are listed in Table 1. Strains VC398 *kin-20(ok505)*X outcrossed x1, MT2257 *lin-42(n1089)*II outcrossed x0, and RB1843 *lin-42(ok2385)*II outcrossed x0 were provided by the Caenorhabditis Genetics Center (CGC), University of Minnesota, USA. Strains RG1590 *lin-42(ox461)*II outcrossed x3 and plasmids pCP2 [*plin-42b/c::isoformb::gfp:unc-54*], pMJ13 [*plin-42b/c::lin42c::gfp::unc54*] and pHG82 [*plin-42a::lin42a::gfp::unc54*] were donated by the Ann E. Rougvie Lab [15]. Strains EG5202 *oxIs12* [*punc-47::gfp, lin-15(+)*] *kin-20(ox423)*X, and EG9581 *oxSi1087* [*pkin-20::rfp::kin-20::kin-20 3’UTR, Cb unc-119(+)* ∗*ttTi5605*] II ; *oxIs12* [*punc-47::gfp, lin-15(+)*] *kin-20(ox423)*X were provided by Erik M. Jorgensen Lab [22], and were later crossed with the VQ1310 *qvIs8* strain, generating the DG1 and DG2 strains, respectively. Strain VQ1850 *kin-20*(*syb4198*[*kin-20b/c::mkate-2^degron::HA*])X was constructed by Sunybiotech (Fuzhou, China). The strains JDW136 *lin-42*(*wrd35*[*lin-42::GFP^degron::3xFLAG*])II, JDW233 [wrdSi46*SCMp*::TIR1::F2A::BFP::AID*::NLS::tbb-2 3’UTR*]I, DV3805 [eSi7(*rgef1p::TIR1::F2A::mTagBFP2::NLS::AID::tbb-2 3’UTR*]I were generated as described [39].

Transgenic animals were generated by standard microinjection techniques [55] and the integration of *psur5::luc::gfp* was induced by UV radiation to generate *qvIs8* [56]. Subsequently, 7 crosses were performed to clean the strain of unwanted mutations.

### Molecular constructs

To obtain the promoter of the gene *kin-20*, 2.6 kb upstream of the start codon of the isoform b of the gene *kin-20* were amplified by PCR from fosmid WRM0617dH07 (Supplementary Table 2). A 1045 bp DNA fragment corresponding to the isoform b of *kin-20* was obtained by PCR from genomic DNA of wild-type nematodes (Supplementary Table 2). Subsequently, the fragment was digested with the restriction enzymes XbaI and KpnI (NEB) and cloned into vector *punc-122::dsRed* (*coel::RFP* Plasmid #8938, Addgene) to generate *punc-122::isoform-b::dsRed* (6.8 kb). The *kin-20* promoter (2647 bp) was cloned into the previously generated plasmid (*punc-122::isoform-b::rfp*) by digestion with restriction enzymes SphI and XbaI to generate the *pkin-20::isoform b::rfp* (7887 bp) construct.

### Luminescence assays

For all assays, nematodes were maintained in LD/CW 12:12h. Adult nematodes were treated with the alkaline hypochlorite solution method to obtain synchronized populations of embryos [57]. The harvested embryos were cultured overnight in a 50 mL Erlenmeyer flask with 3.5 mL of M9 buffer (42 mM Na_2_HPO_4_, 22mM KH_2_PO_4_, 85.5 mM NaCl, 1 mM MgSO_4_), 1X antibiotic-antimycotic (Thermo Fisher Scientific) and 10 μg/mL of tobramycin (Tobrabiotic, Denver Farma) at 110 rpm in LD/CW (∼150 / 0 µmol/m^2^·s); 18.5/20°C, Δ = 1.5°C ± 0.5°C) conditions. The next day, L1 larvae were placed on a Petri dish with NGM (0.3% NaCl, 0.25% peptone, 5 μg/mL cholesterol, 1 mM CaCl_2_, 1 mM MgSO_4_, and 1.7% agar in 25 mM potassium phosphate buffer, pH 6.0) and bacterial lawns of *Escherichia coli* strain HB101.

Two days later, L4 stage nematodes were picked onto 96-well plates (50nematodes/well). They were visualized on a SMZ100 stereomicroscope equipped with an epi-fluorescence attachment (Nikon) with a cool Multi-TK-LED light source (Tolket) to avoid warming of the plate. They were washed once with M9 buffer to remove all traces of bacteria and resuspended in 200 μL of luminescence medium. Luminescence medium contained Leibovitz’s L-15 media without phenol red (Thermo Fisher Scientific) supplemented with 1X antibiotic–antimycotic (Thermo Fisher Scientific), 40 μM of 5-fluoro-2′-deoxyuridine (FUdR) to avoid new eclosions, 5 mg/mL of cholesterol, 10 μg/mL of tobramycin (Tobrabiotic, Denver Farma), 1 mM of D-luciferin (Gold Biotechnology) and 0.05% Triton X-100 to increase cuticle permeabilization. All chemical compounds were bought from Sigma-Aldrich (St. Louis, MO) unless otherwise specified.

For the luminescence assays in LD/CW and FR, luminescence from nematode populations (50 nematodes) was measured using a Berthold Centro LB 960 microplate luminometer (Berthold Technologies) stationed inside an incubator (INGELAB) to allow tight control of the light and temperature in each experiment. Microwin 2000 software version 4.43 (Mikrotek-Laborsysteme) was programmed to leave the plate outside the luminometer after each recording to expose nematodes to the environmental cues. The luminescence of each well was integrated for 10 s every 30 min. The scheme used for all experiments were 3 days at a 12 /12 h LD/CW cycle (∼150 µmol/m^2^/s / 0; 15.5/17°C; ZT0, lights on and onset of the cold-temperature phase) and 4 days at the FR condition (DD/WW, dark/17°C). This allows us to do long recordings (8-10 days) and analyze synchronization and real entrainment. In the assays only in FR conditions, we measured the luminescence from nematode populations (50 nematodes per well), with an AB-2550 Kronos Dio luminometer. At ZT12, the plates were transferred to the Kronos luminometer and monitored in DD/WW (dark/20°C, minimal temperature permitted in this luminometer) for 7 days. Recordings from nematode populations were taken every 30 min with an integration time of 1 min.

### Data acquisition and analysis

Luminescence was sampled at 30 min intervals. Background noise was extracted from the raw data obtained from the luminometer. In all cases, the first 12 to 24 hours of recording were removed due to accumulation of the luciferase enzyme. All raw data were analyzed using a Shiny app developed in the laboratory (https://ispiousas.shinyapps.io/circaluc/). The raw data were detrended, smoothed and normalized to the initial maximum value of each sample and plotted using R (R Core Team, 2021). All data are shown as mean ± SD or SEM of luminescence as indicated in the figures. In each case, the mean corresponds to a population of nematodes (50 nematodes). Subsequently, the circadian period was calculated from the population data using the Lomb-Scargle (LS) periodogram using the lomb R package [58], within a range of 18 to 37 h and with an oversampling of 30. The acrophase (time at peak) and amplitude of each signal was estimated using the Cosinor method, by fitting a cosine waveform to the data using a non-linear least squares regression implemented using the nls function of base R and obtaining the R2 of the fit. Any signal resulting from population analysis with a 24 h period and an R2 adjustment ≥ 0.5 was considered “Synchronized” under cyclic training conditions. In the case of free running rhythms, any signal resulting from the analysis with a period range between 18 h and 38 h, and an R2 adjustment ≥ 0.5 was considered “Circadian”. For statistical analyses, the GraphPad Prism 7 software was used. Statistical significance was set at alpha = 0.05. Final figures were generated using Biorender (https://app.biorender.com/).

### Fluorescence microscopy assays

Nematodes were maintained under cyclic conditions of light and dark and temperature (LD/CW, 12:12h) as described in the strains and maintenance section. For the co-expression analysis of LIN-42 and KIN-20 proteins, nematodes in the L4 stage/young adults were taken from the NGM plate with bacteria and washed twice with M9 buffer. The animals were mounted on a 5% agarose pad and paralyzed with 2% azide. All images were taken using a Leica laser-scanning spectral confocal microscope TCS SP8 (Leica), with a scale of 40X. Lasers to excite GFP, mKate2 and BFP, respectively, were used and two independent experiments were performed with at least 10 nematodes. Image preparation and co-expression analyses were done using ImageJ.

### Protein depletion using the auxin-inducible degron (AID) system

Nematodes were maintained under the same conditions as explained in the luminescence assay section. We added 4 mM of the auxin K-NAA (Phyto-Technology Laboratories, N610) [42] into the luminescent medium on the first day of the assays (ZT=12, Time=0). Control for experiments were performed using L15 (Leibovitz’s L-15 Medium, vehicle) with an equivalent volume. To check the prolonged effect of auxin, BFP positive cells were measured in nematodes exposed to the drug or the vehicle control for 7 days. L4 stage nematodes were used to observe neurons, and L2/L3 stage nematodes were used to observe seam cells. Animals were mounted on a 5% agarose pad and immobilized with 2% azide. All images were taken with a Leica TCS SP8 laser scanning spectral confocal microscope, with a 40X scale. Two independent experiments were performed with at least 10 nematodes each.

### Expression and purification of recombinant protein

The N-terminus of LIN-42 (residues 1-315 or 41-315) was expressed from a pET22-based vector in *Escherichia coli* Rosetta2 (DE3) cells based on the Parallel vector series [59]. The protein was expressed downstream of an N-terminal TEV-cleavable HisGβ1 tag that leaves the sequence ‘GAMDPEF’ on the N-terminus of LIN-42 after TEV cleavage. Cells were grown in LB media at 37°C until the O.D._600_ reached ∼0.8 and then protein expression was induced with 0.5 mM IPTG and cultures were grown for an additional ∼18 h at 18°C. Cells were centrifuged, resuspended in 50 mM Tris, pH 7.5, 300 mM NaCl, 20 mM imidazole, 5% glycerol, 1 mM tris(2-carboxyethyl)phosphine (TCEP), and 0.05% Tween-20. For purification, cells were lysed with a microfluidizer followed by sonication and the lysate was clarified via centrifugation. Ni-NTA affinity chromatography was used to extract HisGβ1-LIN-42 from the lysate and the HisGβ1 was cleaved using His-TEV protease overnight at 4°C in a low imidazole buffer. The cleaved protein was then separated from tag and TEV by a second Ni-NTA affinity column and further purified using size exclusion chromatography on a HiLoad 16/600 Superdex 75 prep grade column (GE Healthcare) in 50 mM Tris, pH 7.5, 200 mM NaCl, 1 mM EDTA, 5% glycerol, 1 mM TCEP, and 0.05% Tween-20. Small aliquots of protein were frozen in liquid nitrogen and stored at −70°C.

### Limited proteolysis and mass spectrometry of recombinant protein

Limited proteolysis of purified protein was performed at 1.5 mg/mL in 25 mM HEPES pH 7.5, 200 mM NaCl, 5 mM DTT with sequencing-grade trypsin (Promega) for one hour at room temperature with 1:800 or 1:1600 mass (w/w) ratios with trypsin for the indicated timepoints. Reactions were quenched with addition of an equal volume of 2X SDS Laemmli buffer (Bio-Rad) and samples were boiled at 95°C for 5 minutes. Digested fragments were resolved by 20% SDS-PAGE and visualized by Coomassie stain. Samples for mass spectrometry were quenched by addition of formic acid to a final concentration of 1% (v/v).

Samples were desalted and separated by HPLC (Surveyor, Thermo Finnegan) on a Proto 300 C4 reverse-phase column with 100 mm x 2.1 mm inner diameter and 5 μm particle size (Higgins Analytical, Inc) using a mobile phase consisting of Solvent A: 0.1% formic acid in HPLC grade water and Solvent B: 0.1% formic acid in acetonitrile. The samples were analyzed on an LTQ Orbitrap linear ion trap mass spectrometer system (Thermo Finnegan). Proteins were detected by full scan MS mode (over the m/z 300-2000) in positive mode with electrospray voltage set to 5 kV. Mass measurements of deconvoluted ESI mass spectra of the reversed-phase peaks were generated by Magtran software.

### Crystallization and structure determination

Crystallization was performed by hanging-drop vapor-diffusion method at 22°C by mixing an equal volume of LIN-42 N-terminus residues 41-315 (4 mg/mL) with a reservoir solution of 150 mM lithium sulfate and 1.5 M sodium potassium tartrate. Hanging drops were also seeded using a cat whisker dipped in crushed LIN-42 crystals obtained from the conditions above, but in a protein:reservoir ratio of 2:1. The crystals were looped and briefly soaked in a drop of reservoir solution with 20% glycerol as a cryopreservant and then flash-cooled in liquid nitrogen for X-ray diffraction data collection. Data sets were collected at the APS beamline 23-ID-D. Data were indexed, integrated and merged using the CCP4 software suite [60]. Structures were determined by molecular replacement with Phaser MR [61] using the mouse PER2 PAS-B domain (PDB: 3GDI). Model building was performed with Coot [62] and structure refinement was performed with PHENIX [63]. All structural models and alignments were generated using PyMOL Molecular Graphics System 2.5.1 (Schrödinger, LLC).

## Acknowledgments

This work was supported by scholarships to M.L.L. from CONICET and from the Emerging Leaders in the Americas Program from Canada, as well as by grants from the Agencia Nacional de Promoción de la Investigación, el Desarrollo Tecnológico y la Innovación (FONCyT, Argentina, to D.G.), the Universidad Nacional de Quilmes (UNQ, Argentina, to D.G.), the Natural Sciences and Engineering Research Council of Canada RGPIN-2017-06553 (to C.Y.B.) and the US National Institutes of Health GM107069 and GM141849 (to C.L.P.). C.Y.B. is a Fonds de recherche du Québec-Santé Research Scholar. National Institutes of Health (NIH) National Institute of General Medical Sciences (NIGMS) [R01GM138701] to J.D.W. We thank Erik Jorgensen for sharing kin-20(ox423) strains, Ann Rougvie for sharing plasmids encoding lin-42, the Caenorhabditis Genetics Center (CGC), which is funded by NIH Office of Research Infrastructure Programs (P40 OD010440) for some strains and WormBase for its valuable genomic resources.

## Supplementary figure legends

**S1. Structural characterization of the LIN-42 N-terminus. A.** Limited trypsin proteolysis of LIN-42 residues 1-315 at the indicated mass ratios (trypsin:protein) for the indicated timepoints. **B.** Liquid chromatography/mass spectrometry analysis of limited trypsin proteolysis of LIN-42 with calculated masses of the corresponding fragments. **C.** Secondary structure prediction of LIN-42 residues 1-315 (JPred) with domain predictions. **D.** Structural alignment of the LIN-42 PAS-B (PDB 8GCI, gray) with the AlphaFold model (https://alphafold.ebi.ac.uk/entry/Q65ZG8) colored by the model confidence. **E.** Predicted aligned error of the LIN-42 AlphaFold model. The shade of green at position (x, y) indicates the expected position error at residue x when the predicted and true structures are aligned on residue y.

**S2. Circadian rhythms occur in *lin-42* mutants under FR.** Representative single traces of luciferase activity rhythms of adult populations shown in control (**A**), *lin-42(ox461)* (**B**), *lin-42(n1089)* (**C**), *lin-42(ok2385)* (**D**), under FR conditions.

**S3. *lin-42* mutants are rhythmic under cyclic and FR conditions.** Representative luciferase activity rhythms of adult populations under dual cyclic conditions (LD/CW, ∼150/0 µmol/m^2^/s; 15.5°C/17°C) and FR conditions (DD, 17°C): control (**A,** n=37), *lin-42(ox461)* (**B**, n=32), *lin-42b* overexpression (OE) transgene strain (**C**, n=20). Luminescence signals are shown as mean ± SEM in red line and all the individual wells are represented in the gray line. Each population consisted of 50 adult nematodes per well. The analysis includes three biological replicates for each strain.

**S4. Circadian rhythms in *lin-42(n1089)* and *lin-42(ok2385)* strains. A**-**C**. Representative luciferase activity rhythms of adult populations under dual cyclic conditions (LD/CW, ∼150/0 µmol/m^2^/s; 15.5°C/17°C) and FR conditions (DD, 17°C), control (**A**), *n1089* mutants (**B**) and *ok2385* mutants (**C**). Luminescence signals are shown as mean ± SEM. The average reported activity was displayed with populations showing a similar first peak in FR conditions. Each population consisted of 50 adult nematodes per well. **D**-**F.** Average reporter activity of rhythmic adult populations under dual cyclic conditions and FR conditions, control (**D**, n=37), *n1089* mutants (**E**, n=40) and *ok2385* mutants (**F**, n=30). Luminescence signals are shown as mean ± SEM in red line and all the individual wells are represented in the gray line. The analysis includes three biological replicates for each strain. **G.** Representation of the acrophase distribution in Rayleigh plots under dual cyclic conditions (LD/CW, blue dots) and FR (DD/WW, red dots) for rhythmic population nematodes: control (LD/CW: 14.55 ± 0.27 h, n=37; R=0.90 and DD/WW: 7.85 ± 0.85 h, n=37; R=0.07), *lin-42(ok2385)* mutants (LD/CW: 15.24 ± 0.17 h, n=30; R=0.96 and DD/WW: 8.63 ± 0.87 h, n=30; R=0.20) and *lin-42(n1089)* mutants (LD/CW: 14.79 ± 0.20 h, n=40; R=0.94 and DD/WW: 7.52 ± 0.72 h, n=40; R=0.27). Arrows represent the average peak phase of *sur-5::luc* expression (mean vectors for the circular distributions) of each group. The length of the vector represents the strength of the phase clustering while the angle of the vector represents the mean phase. Individual data points are plotted outside the circle. The central circle represents the threshold for p=0.05. **H.** Average endogenous period of *lin-42* mutants vs control: control (24.4 ± 0.56 h, n=37), *n1089* mutants (25.34 ± 0.67 h, n=40), *ok2385* mutants (25.91 ± 0.60 h, n=30). One-way ANOVA followed by Dunnett’s multiple comparisons test, ns, p>0.05.

**S5. Circadian rhythms occur in *kin-20* mutants under FR.** Representative single traces of luciferase activity rhythms of adult populations shown in control (**A**), *kin-20(ok505)* (**B**), *kin-20(0×423)* (**C**), under FR conditions.

**S6. *kin-20* mutants are rhythmic under cyclic and FR conditions.** Representative luciferase activity rhythms of adult populations under dual cyclic conditions (LD/CW, ∼150/0 µmol/m^2^/s; 15.5°C/17°C) and FR conditions (DD, 17°C): control (**A,** n=37), *lin-42(ox461)* (**B**, n=32), *lin-42b* overexpression (OE) transgene strain (**C**, n=20). Luminescence signals are shown as mean ± SEM in red line and all the individual wells are represented in the gray line. Each population consisted of 50 adult nematodes per well. The analysis includes three biological replicates for each strain.

**S7. Co-expression of LIN-42B and KIN-20B in neurons and seam cells. A**. LIN-42B::GFP (green channel) and KIN-20B::mKate2 (red channel) are detected in the neurons which also express BFP (blue channel, white circle) in L4 nematodes. Scale bars represent 20 µm. **B.** LIN-42B::GFP (green channel) and KIN-20B::mKate2 (red channel) are detected in the seam cells which also express BFP (blue channel, arrowheads) in L3/L4 nematodes. Scale bars represent 20 µm. **C.** Representative expression of *lin-42* (green bars) and *kin-20* (orange bars) in pharyngeal neurons, motor neurons, sensory neurons, and interneurons. Data obtained from CeNGEN (https://www.cengen.org/). The scale represents the percentage of expression of the genes of interest. Each bar represents a neuron.

**S8. Expression of BFP in neurons and seam cells exposed to Auxin.** Representative images of nematodes expressing the neuronal rgef-1p::TIR-1::F2A::BFP::AID*::NLS::tbb-2, (**A**) and the seam cell SCMp::TIR-1::F2A::BFP::AID*::NLS::tbb-2 reporters (**B**), treated with vehicle and 4 mM K-NAA for 7 days. An overlay of DIC and BFP images was used to show the expression of BFP-positive neuronal cells in animals at stage L3/L4 (arrowheads). Scale bars represent 20 µm.

**S9. A.** Average endogenous period of control (25.09 ± 0.68 h, n=29), *lin-42b*::AID mutants (27.62 ± 1.14 h, n=13) and *kin-20b*::AID mutants (26.93 ± 1.30 h, n=12), which also express the construct *rgef-1p*::TIR-1::F2A::BFP::AID*::NLS::tbb-2. One-way ANOVA, Dunnett’s multiple comparisons test, ns. **B.** Representative luciferase activity rhythms of adult populations shown in **A**, under dual cyclic conditions (LD/CW, ∼150/0 µmol/m2/s; 15.5°C/17°C) and FR conditions (DD, 17°C). **C.** Average endogenous period of control (25.09 ± 0.68 h, n=29), *lin-42b*::AID mutants (27.87 ± 0.74 h, n=14) and *kin-20b*::AID mutants (26.16 ± 1.12 h, n=17), which also express the construct *SMCp*::TIR-1::F2A::BFP::AID*::NLS::tbb-2. One-way ANOVA followed by Dunnett’s multiple comparisons test, ns. **D.** Representative luciferase activity rhythms of adult populations shown in C, under the same dual cyclic conditions and FR conditions. Each population consisted of 50 adult nematodes per well. The analysis includes three biological replicates for each strain.

**S10. *lin-42*::AID and *kin-20*::AID are rhythmic under cyclic and FR conditions.** Representative luciferase activity rhythms of adult populations under dual cyclic conditions (LD/CW, ∼150/0 µmol/m^2^/s; 15.5°C/17°C) and FR conditions (DD, 17°C) in the control strains, *lin-42*::AID, *kin-20*::AID, *lin-42*::AID;*kin-20*::AID, with vehicle (A/C) and auxin (B/D), in neuronal cells (A-B) and in seam cells (C-D). Luminescence signals are shown as mean ± SEM in red line and all the individual wells are represented in the gray line. Each population consisted of 50 adult nematodes per well. The analysis includes three biological replicates for each strain.

