## Supplementary figures and images for "Regulation of the circadian clock in *C. elegans* by clock gene homologs *kin-20* and *lin-42*"

### Supplemental Figure 1

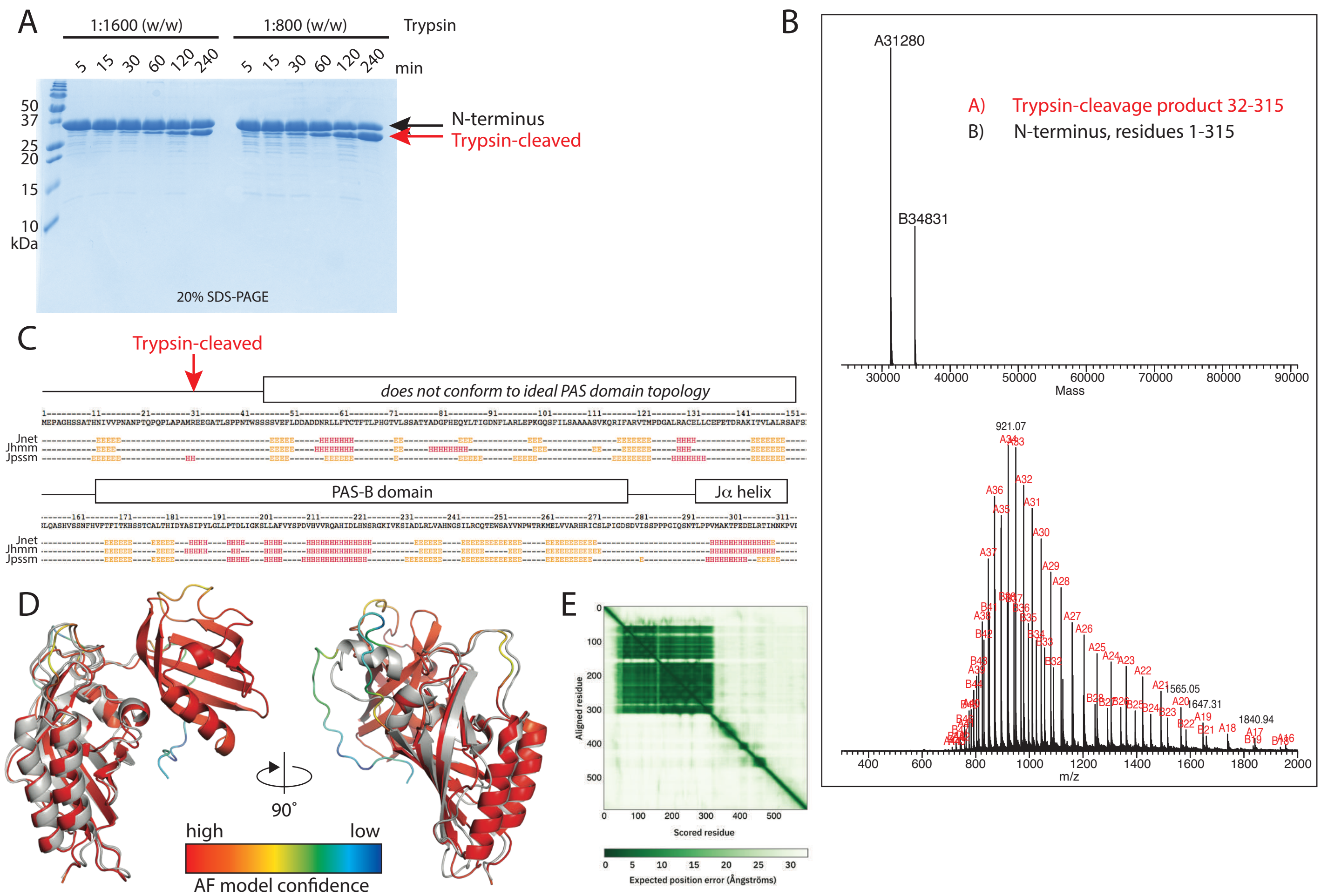

### Supplemental Figure 2

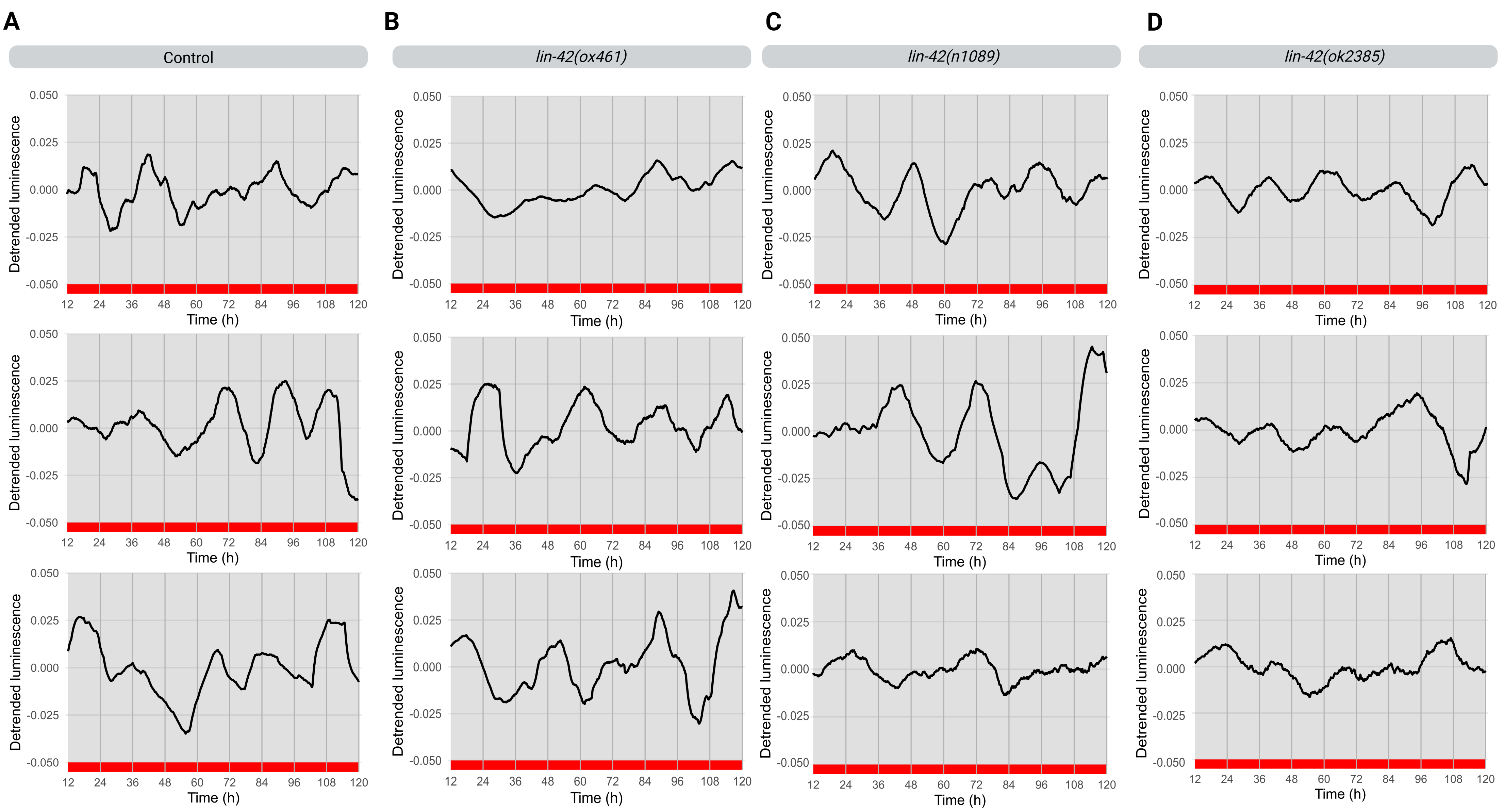

### Supplemental Figure 3

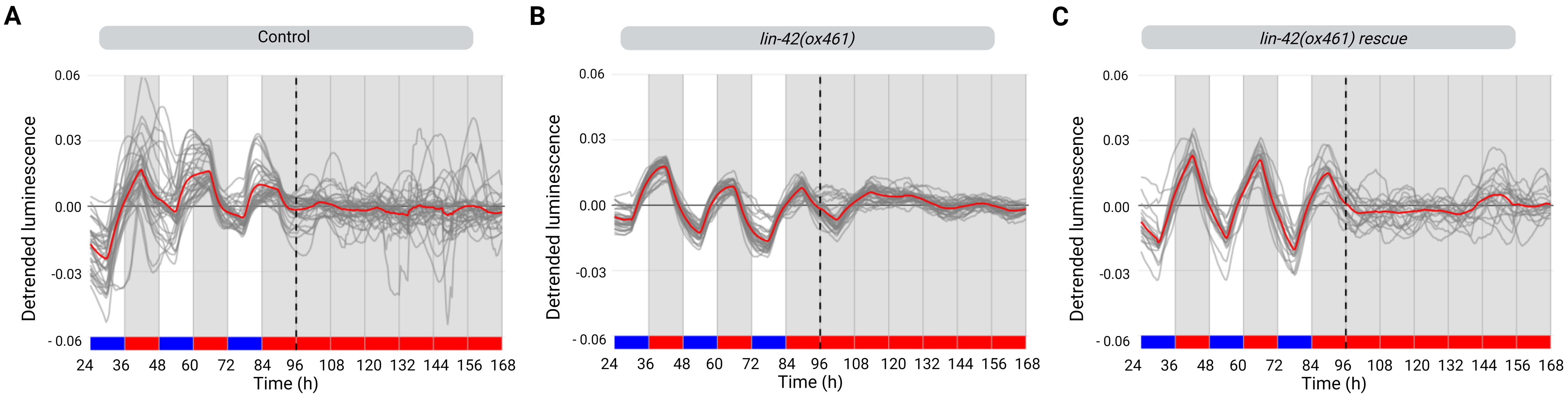

### Supplemental Figure 4

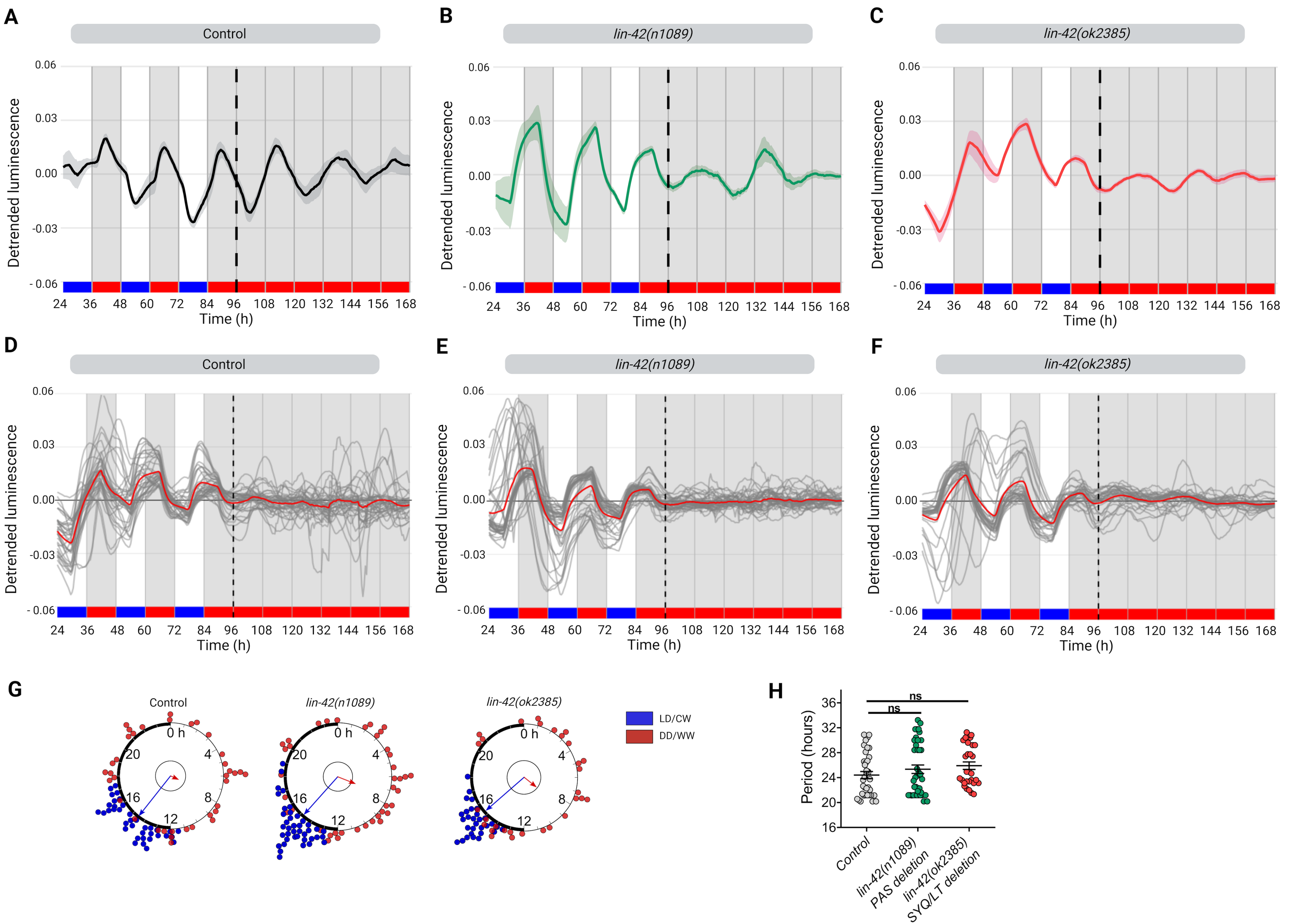

### Supplemental Figure 5

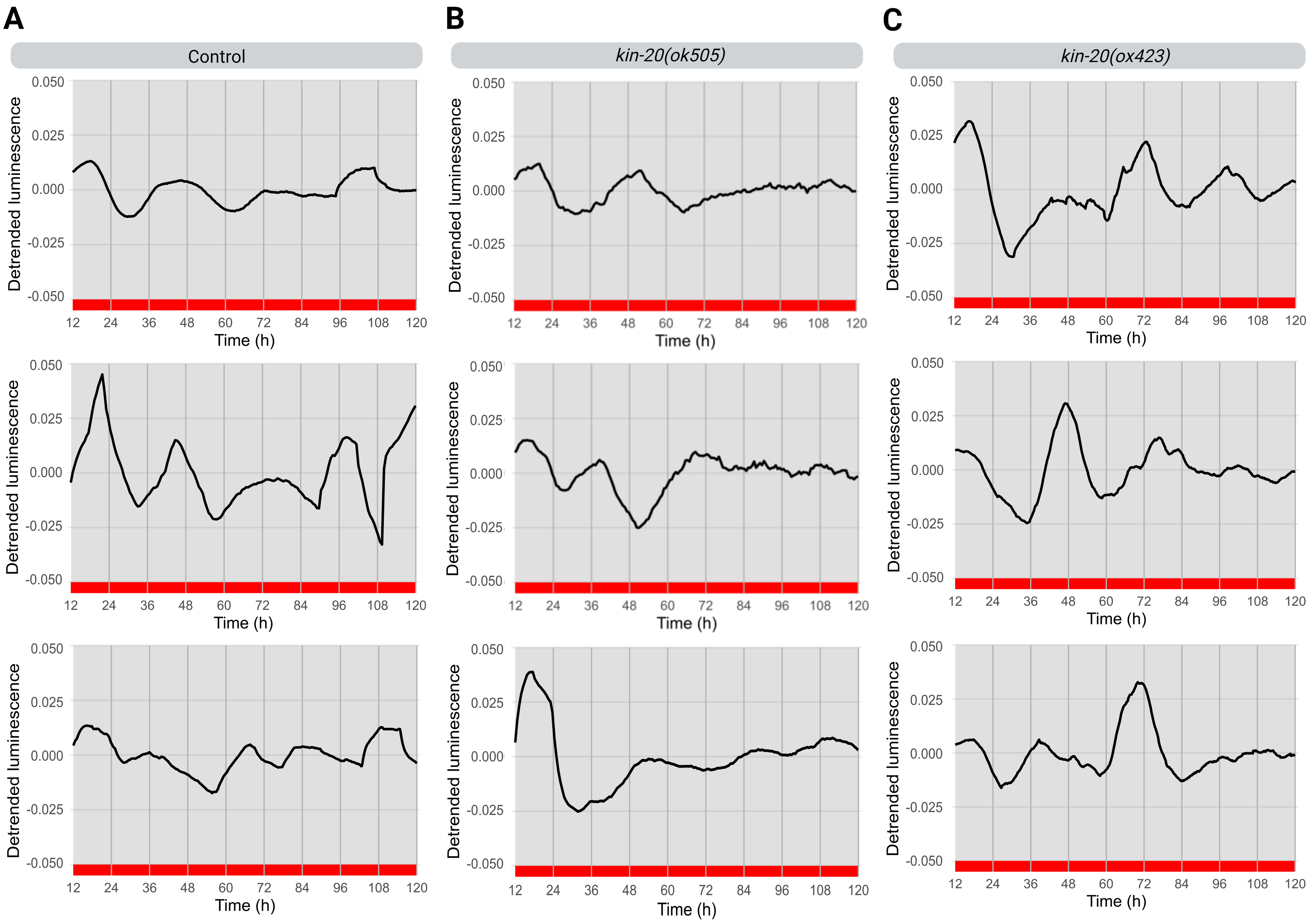

### Supplemental Figure 6

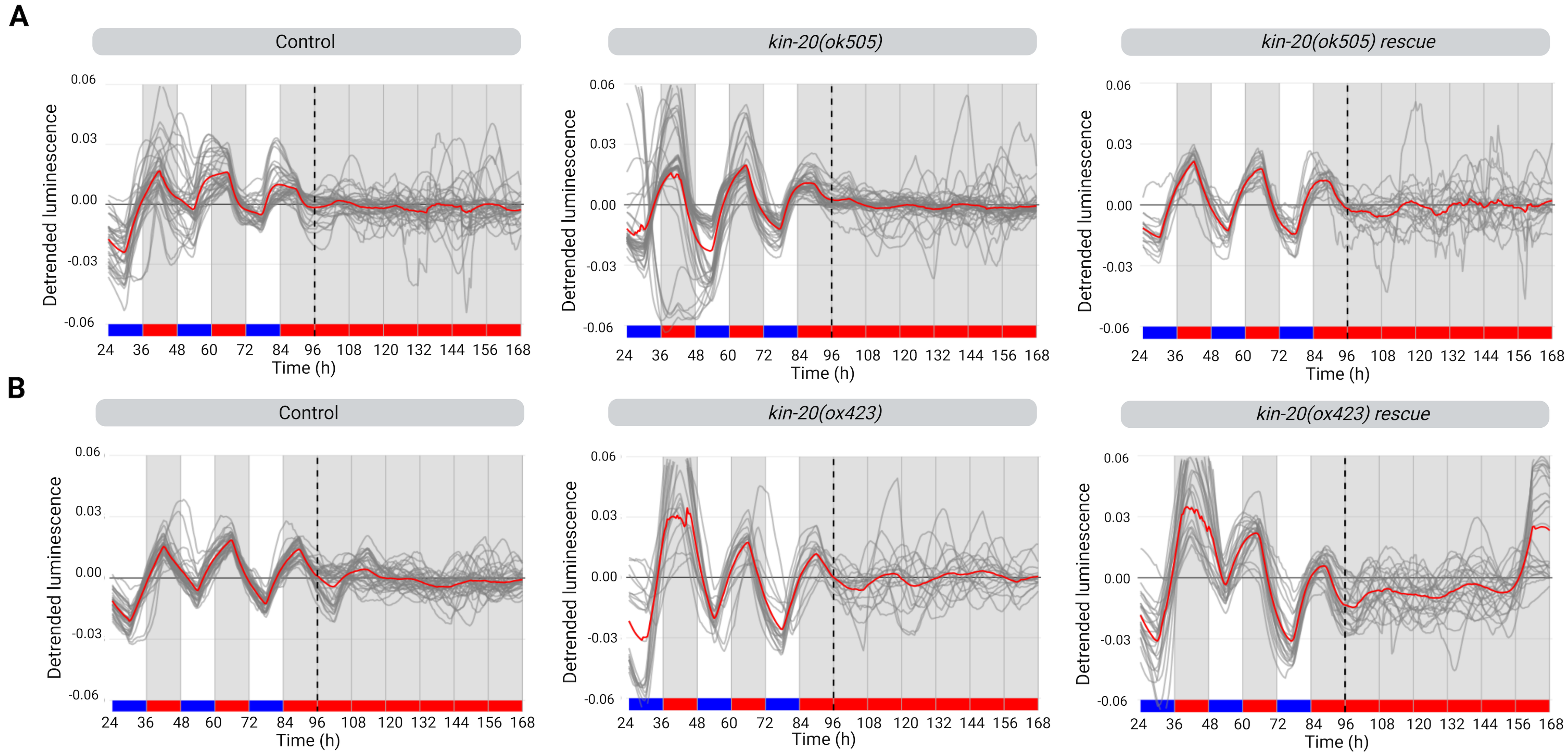

### Supplemental Figure 7

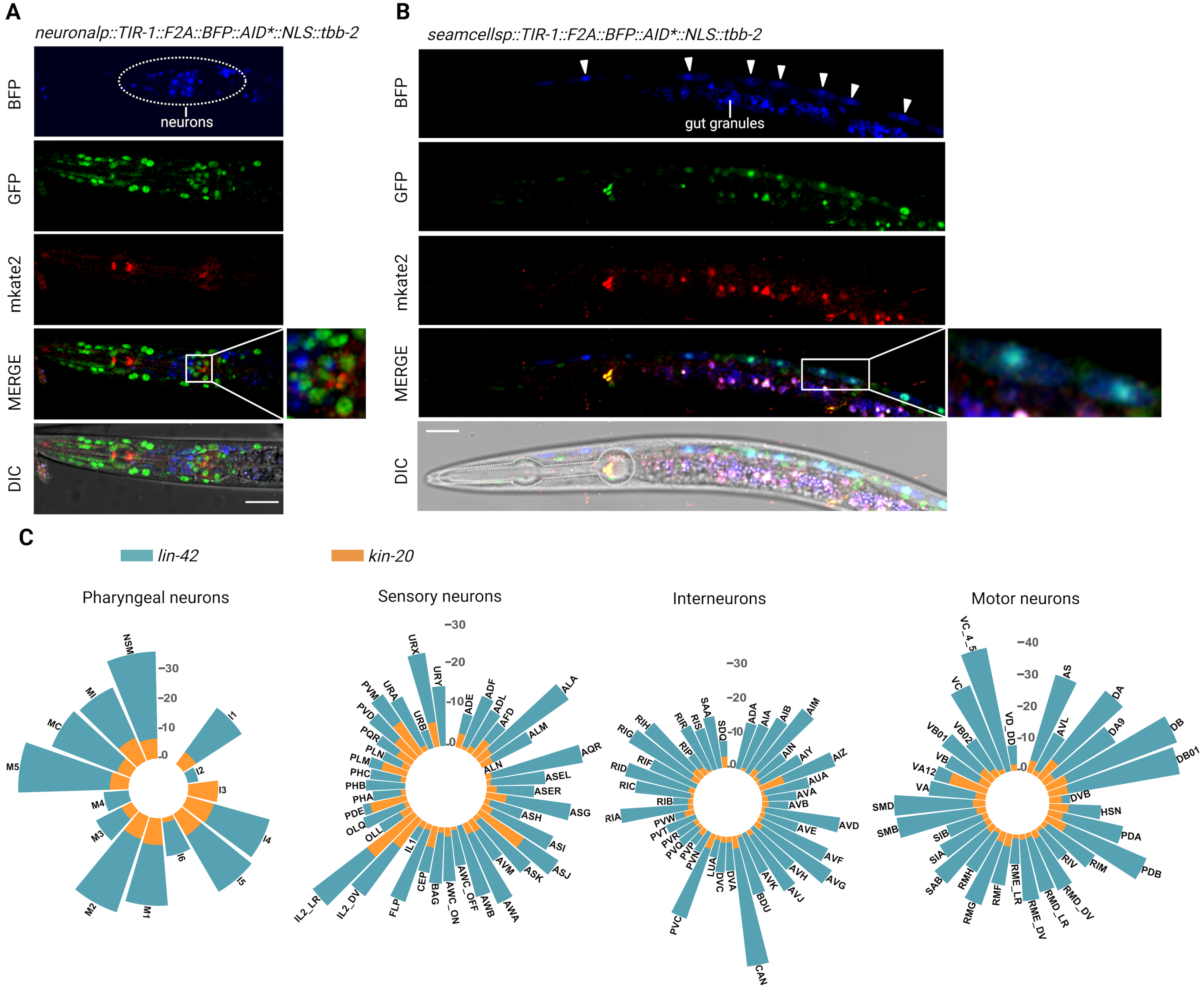

### Supplemental Figure 8

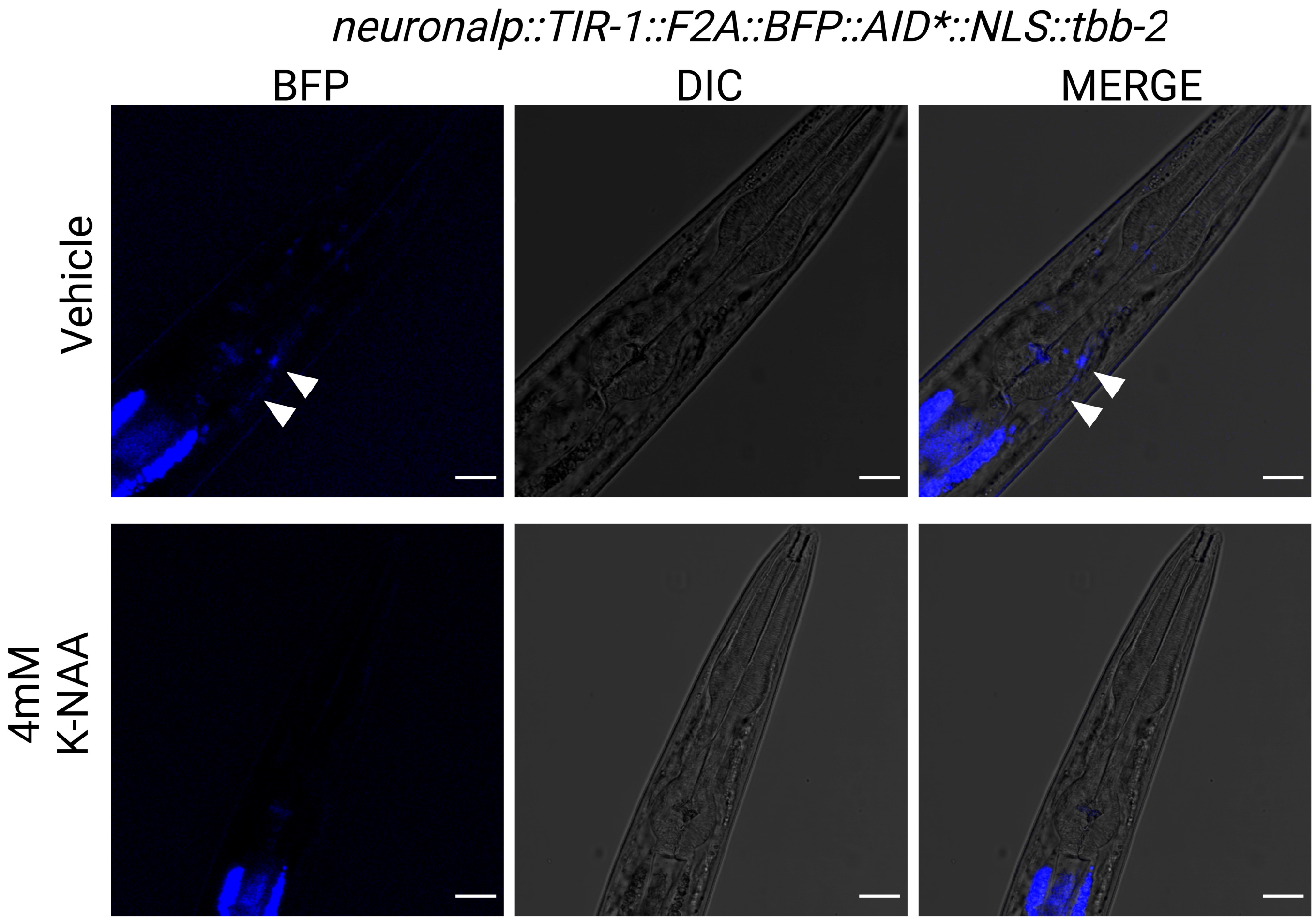

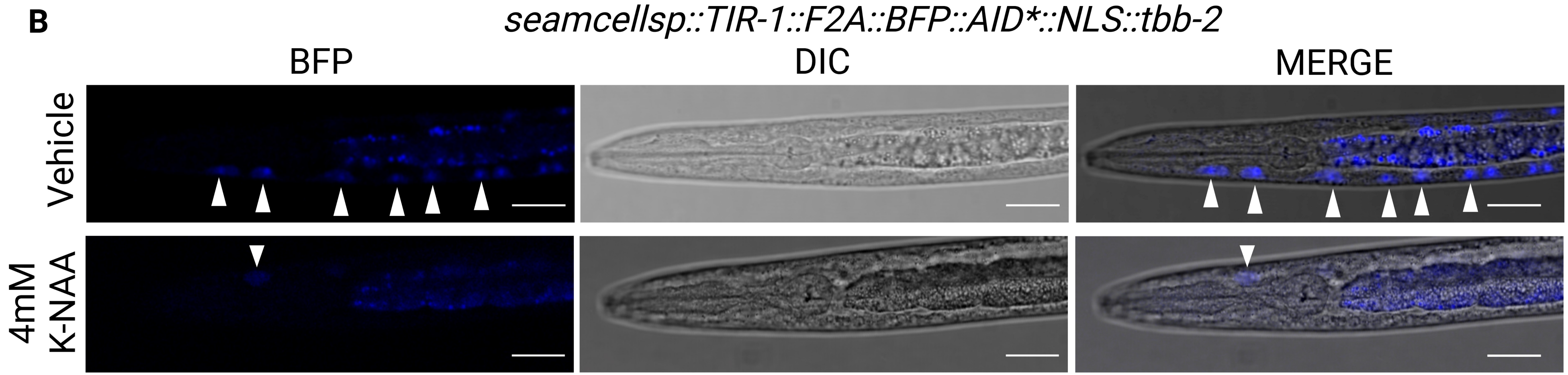

### Supplemental Figure 9

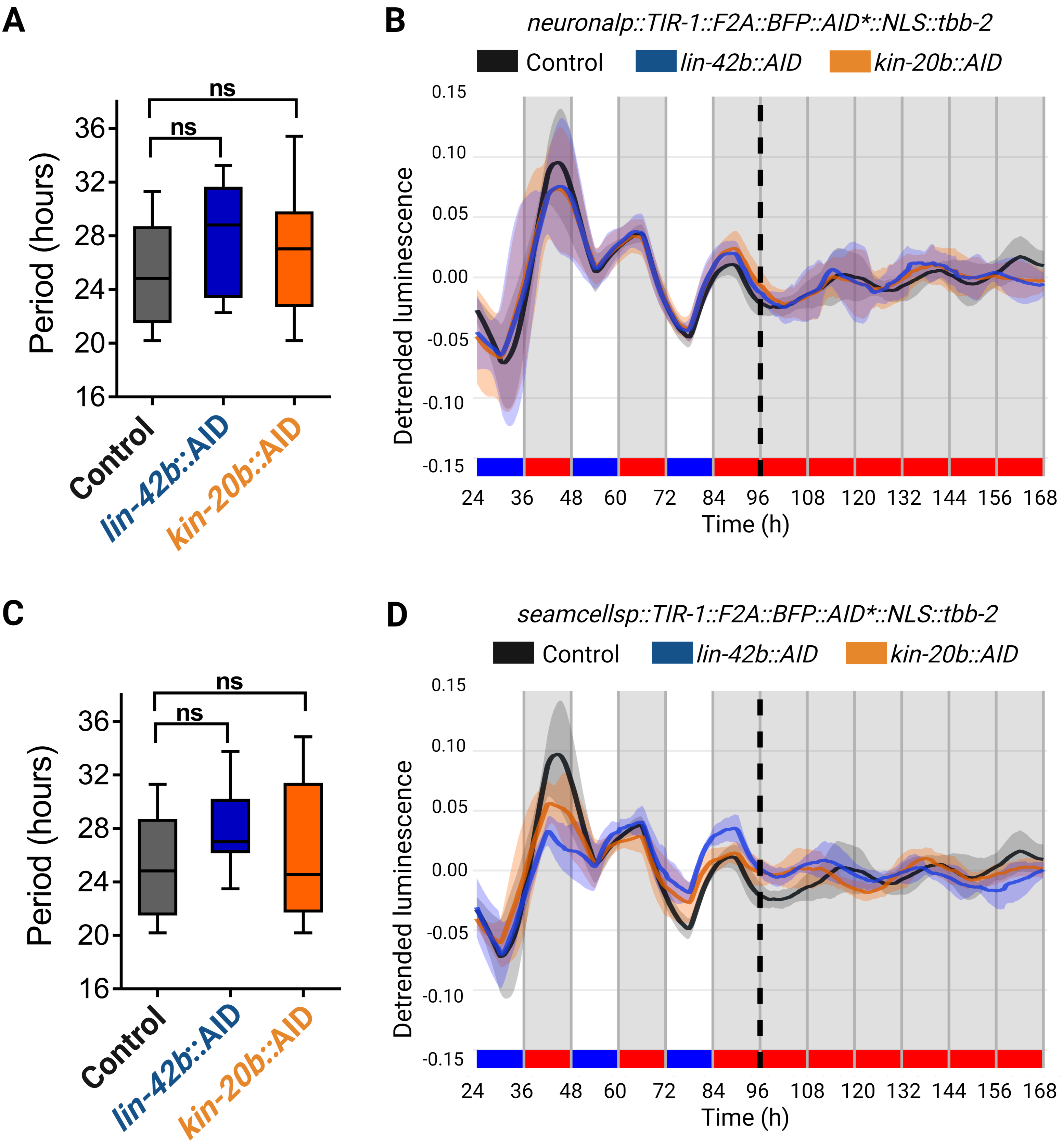

### Supplemental Figure 10

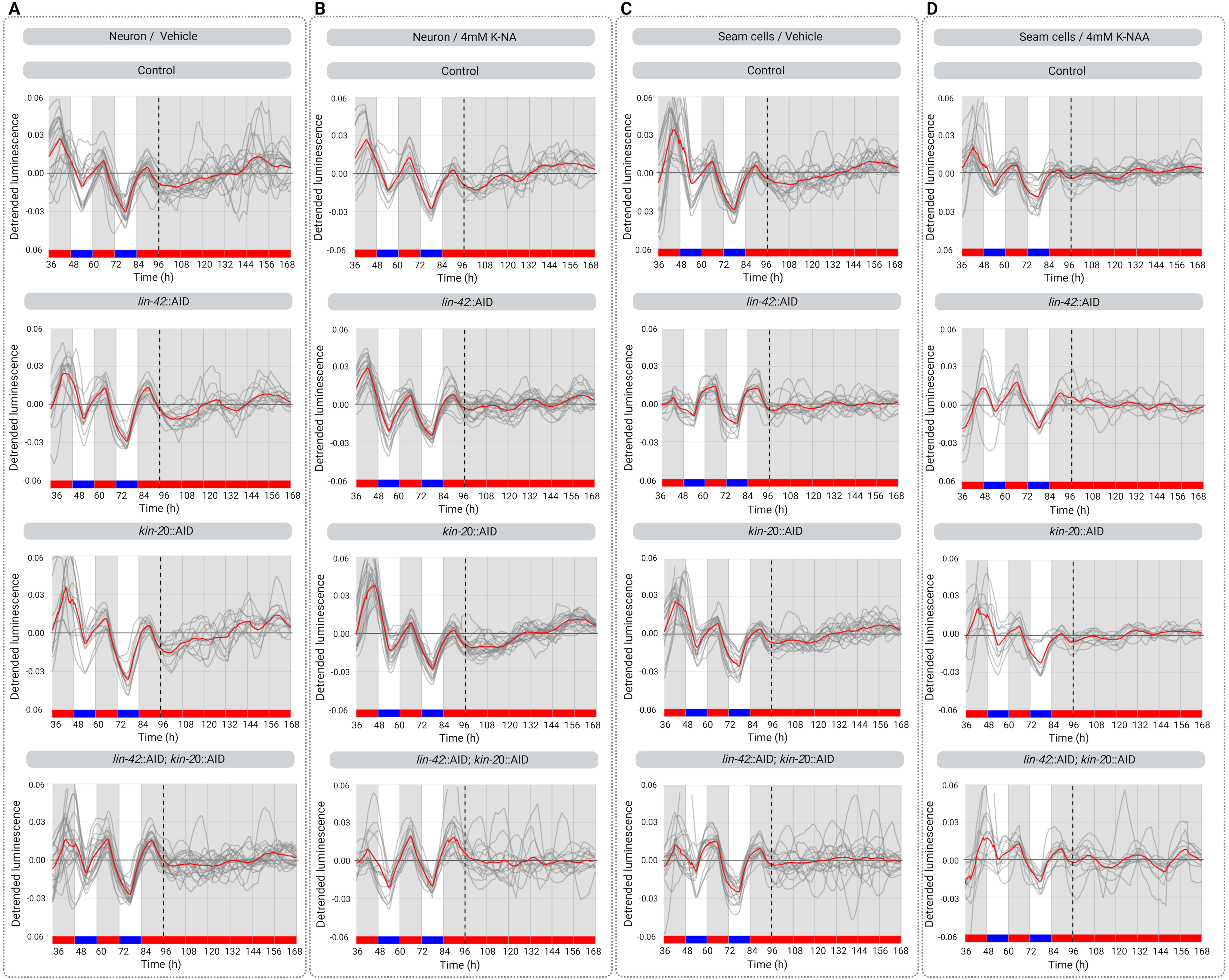
